# Connectivity loss in experimental pond networks leads to biodiversity loss in microbial communities

**DOI:** 10.1101/2024.08.05.606584

**Authors:** Beáta Szabó, Máté Váczy-Földi, Csaba F. Vad, Károly Pálffy, Thu-Hương Huỳnh, Péter Dobosy, Ádám Fierpasz, Zsuzsanna Márton, Tamás Felföldi, Zsófia Horváth

## Abstract

Habitat fragmentation is among the most important global threats to biodiversity, however, the direct effects of its components including connectivity loss are still lesser known. Our understanding of these drivers is especially limited in microbial communities. Here, by conducting a four-month outdoor experiment with artificial pond (mesocosm) metacommunities, we studied the effects of connectivity loss on planktonic prokaryote and microeukaryote communities. Connectivity loss was simulated by stopping the dispersal among local habitats while keeping the habitat amount constant and the abiotic environment homogeneous. We found that connectivity loss led to higher levels of extinction and a decrease in both local and regional diversity in microeukaryotes. In contrast, diversity patterns of prokaryotes remained largely unaffected, with some indications of extinction debt. Connectivity loss also led to lower evenness in microeukaryotes, likely through changes in biotic interactions with zooplankton grazers. Our results imply that connectivity loss can directly translate into species losses in communities and highlight the importance of conserving habitat networks with sufficient dispersal among local habitats.

## Introduction

Habitat loss is one of the primary drivers of global biodiversity decline (Brooks et al., 2002; Hanski, 2011; Pimm, 2008; WWF, 2018) that can be manifested as two major mechanisms. First, it affects habitat networks directly when species disappear due to the sampling effect related to the species–area relationship (Fahrig, 2013). At the same time, the spatial configuration of the remaining habitats may undergo changes, as well (Fahrig, 2017), which involves both the fragmentation of large areas and an increased spatial isolation through the loss of connectivity between habitat patches (Haddad et al., 2015; Horváth et al., 2019). In addition, functional connectivity loss can occur without significant reduction in total habitat amount via decreasing landscape permeability driven by increasing urbanization and land use changes (Oertli and Parris, 2019; Trombulak and Frissell, 2000). While these processes can all have a negative impact on biodiversity, their relative roles still represent a highly debated topic in ecology (Fahrig, 2019; Fletcher et al., 2018; Valente et al., 2023).

Connectivity loss can have several negative impacts on biodiversity. First, it can lead to increased probabilities of local extinction by directly decreasing colonization rates (Fahrig, 2002), the exchange of genes and individuals, and contributing to inbreeding (Frankham, 2006; Hagen et al., 2012; Harrisson et al., 2012; Klinga et al., 2019) and ecological drift resulting from demographic stochasticity (Chase et al., 2020; Ryberg et al., 2012). In addition, it can enhance the importance of further stochastic processes, such as priority effects (Chase, 2003; Woody et al., 2007). Connectivity loss can also indirectly modify density-dependent biotic processes (Hagen et al., 2012; Magrach et al., 2014; Peh et al., 2014), resulting in changes in community structure and richness (Berga et al., 2015; Wardle, 2016; Wardle and Zackrisson, 2005), or functional diversity (Wardle, 2016; Wardle and Zackrisson, 2005), ultimately reflected in altered ecosystem functioning (Gonzalez et al., 2011). Additionally, it can decrease the stability and persistence of populations during environmental change (Hassell et al., 1993; McCann et al., 2005). On the other hand, fragmentation might also have a stabilizing effect on unstable resource-consumer relationships via rescue effects and spatial refuges (Briggs and Hoopes, 2004). Moreover, it can reduce the predation, spread of disturbances (Levin and Carpenter, 2012) and diseases(Gonzalez et al., 1998; Graham et al., 2022; Haddad et al., 2017) (Hess, 1994). This overall highlights the importance of considering biotic interactions as underlying mechanisms of fragmentation effects.

Studies on the effects of connectivity loss on biodiversity have so far mostly focused on continuous habitats undergoing fragmentation, mainly in terrestrial biomes (e.g., Gonzalez et al., 1998; Graham et al., 2022; Haddad et al., 2017). Consequently, our understanding of how connectivity loss affects biodiversity in naturally patchy, insular types of ecosystems, such as small standing waters, remains poorly known. Furthermore, most previous studies on habitat fragmentation were either performed under heterogeneous environmental conditions or did not monitor potential environmental drivers (e.g., Evans et al., 2017; Gibb and Hochuli, 2002; Gibson et al., 2013; Magura et al., 2001), making it difficult to disentangle and determine the direct effect of fragmentation. Micro- and mesocosm experiments can address these limitations as under experimental conditions, it is more feasible to disentangle the pure effect of connectivity loss from other factors, such as habitat size and amount, local environmental variables, or interspecific interactions.

Research on connectivity loss has traditionally focused more on macroorganisms (e.g., Hamer, 2016; Hamer et al., 2023; Johnson et al., 2013; Rösch et al., 2013), while there is a clear knowledge gap with especially scarce experimental data with regard to microorganisms. This might partly stem from the phenomenon that microorganisms have generally higher dispersal rates compared to macroorganisms, and are consequently considered to be less dispersal limited (Baas Becking and Nicolai, 1934; Finlay, 2002; Foissner, 2006). Nevertheless, their role as primary producers, decomposers and as a link in energy transfer to higher trophic levels makes them important members of the food webs and highlights the need for more research on their sensitivity to connectivity loss. Existing experimental studies on spatial connectivity between aquatic habitats, however, have generally targeted single trophic levels. In these studies, organisms representing higher trophic levels, such as zooplankton, were typically investigated in mesocosms (Gianuca et al., 2017; Howeth and Leibold, 2013, 2010; Sinclair and Arnott, 2018; Thompson et al., 2023; Thompson and Shurin, 2012) while organism groups at lower trophic levels (e.g. diatoms and other microalgae) were rather studied in laboratory microcosms (de Boer et al., 2014; Eggers et al., 2012; Guelzow et al., 2017). There have been relatively few studies encompassing multiple trophic levels within the same experimental setup. Furthermore, these usually applied connectivity to an external regional species pool (Limberger et al., 2019; Turunen et al., 2018; Vad et al., 2023). Modeling more realistic networks and starting from setups with within-network connectivity (as in large-scale terrestrial experiments starting with an undisturbed system) could provide setups more powerful to reveal the direct effects of connectivity and fragmentation, and eliminate bias due to potential mass effects produced by the repeated flow of organisms from an external source.

In the current study, we aimed to investigate the effects of habitat network fragmentation (i.e., the loss of connectivity between habitats) on biodiversity in experimental ponds, by keeping habitat amount and abiotic environment constant. More specifically, we tested whether habitat fragmentation in the form of connectivity loss among habitats reduces the local and regional richness of aquatic microorganisms in a four-month long experiment. In this regard, we compared the response of prokaryotes and microeukaryotes, and expected that prokaryotes will be less affected by fragmentation due to their smaller size (De Bie et al., 2012) and larger population size (Berninger et al., 1991; De Bie et al., 2012; Fenchel, 1988). We also explicitly tested whether the local richness of microorganisms might be affected by food web interactions, specifically the biomass of their grazers, zooplankton, and whether the observed relationship is modulated by fragmentation. Ultimately, we aimed to identify the drivers of survival probability, including the direct and indirect effect of fragmentation and the role of initial regional abundance.

## Materials and methods

### Experimental setup and sampling

We created six metacommunities (M1-M6) in an outdoor experimental setup. Each metacommunity consisted of five artificial ponds (mesocosms), with a total of 30 mesocosms set up in a spatially randomized arrangement. As mesocosms, we used 225-L UV-resistant and food-safe PEHD (high-density polyethylene) plastic barrels, filled up to a 200-L experimental volume. They were covered with mosquito nets to exclude macroinvertebrates and prevent coarse organic material (e.g. leaves) from falling into the mesocosms.

The mesocosms were filled with tap water and allowed to stand for de-chlorination by evaporation for four days. Then, each mesocosm was inoculated with the inoculum representing a pooled plankton community from 10 intermittent lowland pools and ponds in the Pannonian Ecoregion (Text S1, Table S1). This inoculum was equally distributed among the mesocosms by replacing 21 L of water taken out from each mesocosm. Thereby, we ensured that the initial communities were identical in all mesocosms, containing microbial communities representative for natural lowland ponds. Nutrient concentrations were adjusted to match those measured in natural ponds, which were used for inoculation (0.7 mg L^-1^ total phosphorus, with 2.1 mg L^-1^ total nitrogen concentrations; Boros et al., 2017; Vad et al., 2017).

The experiment took place between 7^th^ July and 27^th^ October 2020. Within each of the six metacommunities, dispersal treatment by 1% weekly water exchange was performed for 4 weeks (7^th^ July - 4^th^ August; Figure 1). This was carried out by taking out 2.5 L of water with a transparent plexiglass tube plankton sampler from each mesocosm of a metacommunity after thoroughly homogenizing (stirring with a long stick) the water. Then, the pooled water sample of the five mesocosms was mixed and redistributed into the same mesocosms thereby ensuring that 1% of the volume of each mesocosm, i.e., 2 L was exchanged with the other four mesocosms of the metacommunity. After four weeks, we started the fragmentation treatment in three of the six metacommunities. To achieve this, we stopped the weekly dispersal treatment in three metacommunities (hereinafter referred as ‘fragmented metacommunities’, M4-M6) to simulate connectivity loss while it was continued in the three control (hereinafter referred to as connected) metacommunities (M1-M3) for another 12 weeks (until 27^th^ October; Figure 1). To ensure that the disturbance associated with the dispersal treatment did not cause any systematic differences between treatments, water in the fragmented mesocosms was also homogenized each week, however, without water exchange among mesocosms. Sampling equipment was always washed with tap water between mesocosms and experimental dispersal events to avoid cross-contamination.

**Figure 1.**
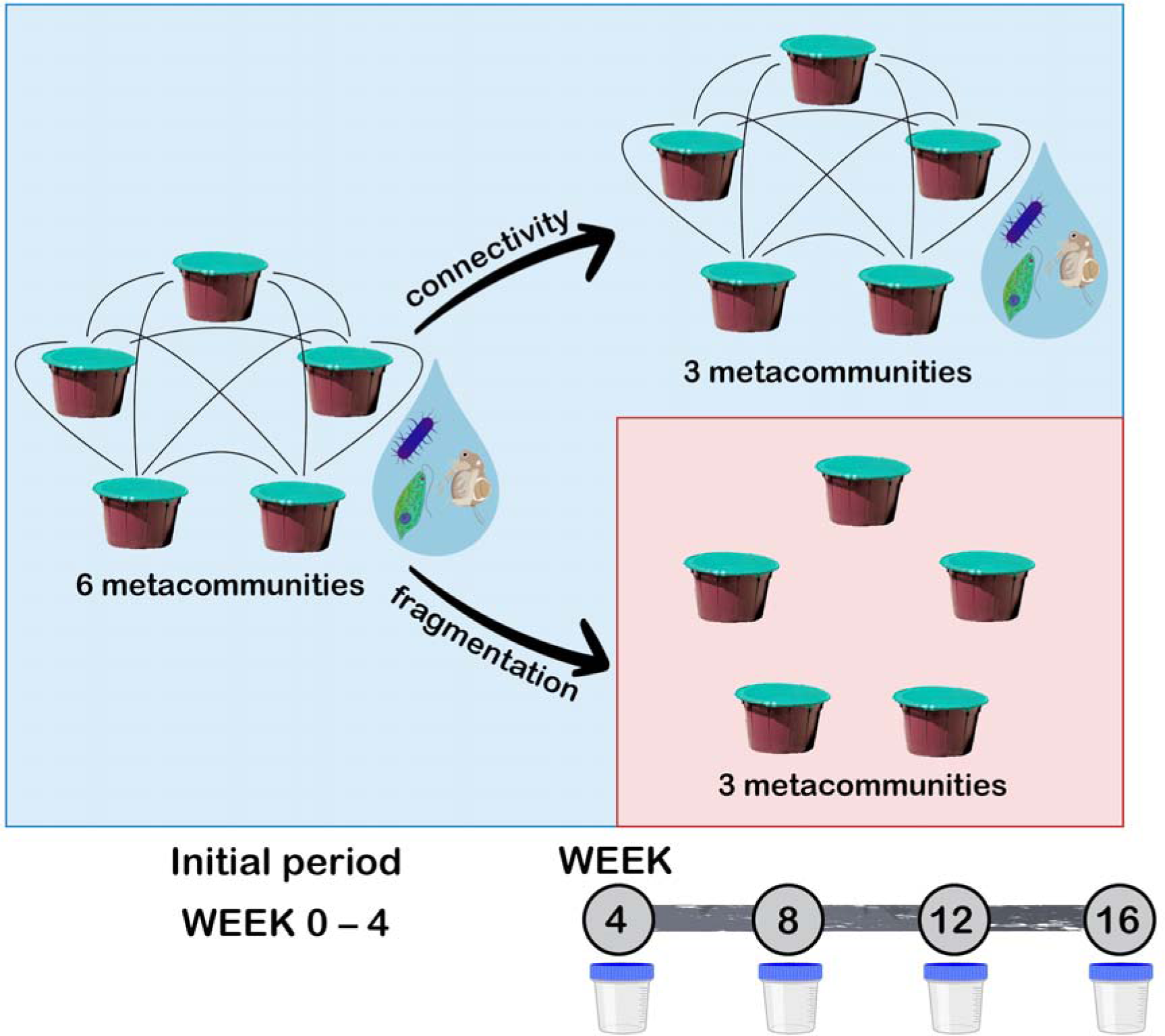
General setup of the experiment. Blue and red panels represent the two types of treatment (connectivity vs. fragmentation) applied for the metacommunities consisting of five habitats. Black solid lines between the habitats illustrate the weekly 1% water exchange. Sampling events are indicated by sample containers.

We sampled all mesocosms at the start of the fragmentation treatment and from that point, every four weeks resulting in four sampling events in total (Figure 1). To ensure a homogeneous distribution of plankton, each mesocosm was stirred prior to sampling and a total of 1.5 L of water was taken out using the same plexiglass tube sampler also used for dispersal. Water was filtered through a 100-μm mesh plankton net to remove most multicellular organisms. 250 mL of the filtered water was taken to the laboratory in a cooling box and filtered through an MF-Millipore membrane filter (Merck, 25 mm Ø, 0.1 μm pore size) to retain material for eDNA-based analyses. The filtered volume depended on the actual microbial biomass as we filtered water until filters were clogged or we reached 250 mL of total volume. The filters were then stored at -20 °C until processing.

To track phytoplankton dynamics, we used chlorophyll-a fluorescence (maximum fluorescence yield, Chl-a, a.u.) as a proxy of total phytoplankton biomass, which was measured with a handheld fluorometer (AquaPen AP 110-C, Photon System Instruments) at each of the four sampling events. Additional samples for the microscopic analysis of phytoplankton communities were taken during the last sampling event (week 16). 50 mL aliquots of the same 1.5 L representative sample that we collected for the molecular samples were fixed with 50 μL Lugol’s iodine and stored at 5 °C until subsequent analysis. Zooplankton sampling was also carried out at each sampling event. To do so, a total of 4 L of water was collected with the plexiglass tube sampler from three randomly chosen spots (1.5 L + 1.5 L + 1 L) of each mesocosm after thorough mixing. Thereafter, the pooled water was filtered through a 30-μm mesh plankton net and zooplankton samples were preserved in 70% ethanol. In parallel, abiotic environmental variables such as temperature, pH, electric conductivity, total phosphorus and total nitrogen were also analyzed (Table S2). For a more detailed description of sample processing of phytoplankton, zooplankton, measurement of chlorophyll-a fluorescence, and determination of physical and chemical variables, see Text S2.

From week 8, filamentous algae appeared at the water surface in the mesocosms. To be able to account for their potential effect on the microbial communities and possible treatment-specific differences in their presence and quantities, we also sampled them in week 8, 12 and 16 (Table S3). Their sampling was carried out from the same 1.5 L water sample that we collected for the molecular samples. Filamentous algae that were present in this sample were retained on a 100-μm mesh plankton net, immediately frozen and their dry weight after freeze-drying was determined later in the lab (referred to as filamentous algae biomass - FB, g L^-1^).

### DNA isolation, amplification, sequencing and data processing

DNA extraction from the material retained on the membrane filters was carried out using the DNeasy PowerSoil Kit (Qiagen, Germany). The isolated DNA was amplified through polymerase chain reaction (PCR) to increase the quantity of fragments of taxonomic marker genes encoding 16S and 18S ribosomal RNA. The PCR products were then sent to Genomics Core Facility RTSF, Michigan State University, USA for sequencing on an Illumina MiSeq platform (Illumina, USA). Raw sequencing reads were processed with mothur v.1.47.0 (Schloss et al., 2009). For further information on the isolation, amplification, sequencing, and data processing procedures, refer to Text S2.

### Statistical analysis

To track whether patch homogeneity lasted throughout the experiment, we tested for any potential treatment-specific differences between abiotic environmental drivers, FB (as competitors), and the taxon richness (αS) and biomass (ZB, µg L^-1^) of zooplankton (as grazers) that might have affected the microbial metacommunities. In the case of environmental variables, permutational multivariate analysis of variance (PERMANOVA, permutations=2000) was performed based on Euclidean distance of z-score standardized data using the nested.npmanova() function in the ‘BiodiversityR’ v. 2.15-4 package (Kindt and Coe, 2005). In the case of abiotic parameters, we did not find any systematic differences among the connected and fragmented metacommunities (Table S2, S4) that could have introduced bias via environmental heterogeneity. Analysis of variance (ANOVA) was run on αS and ZB for each of the four sampling events, and on FB for week 8, 12 and 16. In each PERMANOVA and ANOVA, treatment was involved as a fixed, and metacommunity ID as a nested factor (i.e., nested within the treatment). Zooplankton communities were exclusively composed of Rotifera and Cladocera taxa over the experiment (Figure S1, Table S5). A significant treatment effect on ZB emerged in week 16 (Table S6), hence, we included it as an explanatory variable in the further data analyses in order to explore potential causal relationships with the microbial communities. As we found no systematic treatment-specific difference in any of the other potential drivers (Table S3, S4), we omitted using them in the analytical framework used for microbial metacommunity patterns.

To obtain prokaryote and microeukaryote community datasets, we rarefied both 16S and 18S ASV sets separately to the read number of the sample having the lowest sequence number (13835 reads for the 16S set and 2224 reads for the 18S set). All statistical analyses were carried out for the rarefied 16S (hereinafter referred to as prokaryotes) and 18S (microeukaryotes) community datasets separately using the R (v. 4.2.1) programming language (R Core Team, 2022). Raw sequence reads are available in the European Nucleotide Archive (https://www.ebi.ac.uk/ena/browser/view/PRJEB78363) under reference number PRJEB78363, and the ASV sets with the related list of taxa in Table S7-S10.

Separately for each sampling event (week 4, 8, 12, 16), ANOVA was run to test the possible effect of treatment (connectivity vs. fragmentation) on the number of observed ASVs (αS, i.e. local richness), the effective number of ASVs of PIE (αS_PIE_) and Pielou’s evenness (J, referred to as evenness later on, measures how evenly the number of individuals are distributed among the ASVs) at α-scale, and the Whittaker’s β-diversity (β_S_ = γS/αS, i.e. compositional variation) in each metacommunity. In contrast to αS being sensitive to changes in the number of rare species (ASVs in our case), αS_PIE_ rather indicates the changes in the number of abundant species (McGlinn et al., 2019). In the models, treatment was included as a fixed and metacommunity ID as a nested factor. To compare the number of observed ASVs at γ-scale (γS i.e., regional richness) and the extrapolated richness across connected and fragmented metacommunities at each sampling event, sample-size-based rarefaction and extrapolation approach (Chao et al., 2014) using ‘iNEXT’ v. 3.0.0 package (Hsieh et al., 2020) was applied. The 95% confidence intervals were calculated based on 10 bootstrap replications. To test whether treatment had a significant effect on αS over the entire experimental period, generalized additive models (GAMs) were run with treatment as the main linear predictor, adding time (i.e., the week of sampling) with varying shapes of smooth according to individual metacommunities (k=3 and k=4 for prokaryotes and microeukaryotes, respectively). The models were built using the gam() function in the ‘mgcv’ v. 1.8-42 package (Wood, 2017). Model diagnostics (including selection of k values) were inspected with the appraise() function of ‘gratia’ v. 0.8.1 package (Simpson, 2023).

To assess the possible treatment effect on Chl-a (as an indicator of phytoplankton biomass) over the whole experiment, and phytoplankton diversity at the last sampling event (week 16), nested ANOVAs were performed on Chl-a, and αS, αS_PIE_ and J of phytoplankton the same way as carried out for prokaryotes and microeukaryotes.

Linear mixed-effects models (LMMs) using the ‘lme4’ v. 1.1-35.1 package (Bates et al., 2015) were built to explore the potential effect of the treatment and ZB on αS, αS_PIE_ and J in prokaryotes and microeukaryotes in week 16. In the models, treatment and ZB were included as fixed, while metacommunity ID as a nested random effect factor. In each case, we built two models, one without and another with the interaction of treatment and ZB, then compared them with a chi-square test using the anova() function to select the best fit model. The interaction term only improved the model built for αS in microeukaryotes (χ^2^(1, - 15.6191) = 8.323; P<0.01), therefore, we only kept it in this case.

To assess the relationships (direct and indirect effects) between treatment, ZB, and diversity (αS or J) in prokaryotes and microeukaryotes in week 16, structural equation models (SEMs) were built. Here, we chose to only include αS and not αS_PIE_ given that they both describe local taxonomic richness (as opposed to evenness which entails additional information on dominance patterns). To find the best fitting SEM that describes the processes, we started by building simple SEMs considering both potential causal directions linking zooplankton and microbes (Figure S2 and S3). Each SEM consisted of linear mixed-effects models (LMMs), where treatment was included as a dummy variable, and metacommunity ID as a random effect factor nested within treatment. Bootstrapping (1000 randomizations) was applied to determine the significance of standardized path coefficients. The models were compared and selected based on R^2^ and Akaike information criterion (AIC) values. In microeukaryotes, αS was better predicted when the LMM only included the treatment as an explanatory variable (Figure S3) and not both the treatment and ZB (Figure S2). Contrarily, J was better predicted by the LMM that included both the treatment and ZB as explanatory variables (Figure S2). Accordingly, we built a final complex SEM (using pathways with the highest predictive values in the initial models) where the causal pathway assumed an indirect effect of microeukaryote αS on microeukaryote J via ZB, and where the potential treatment effect was also included as a direct effect. In the case of prokaryotes, the simple SEMs only revealed a significant positive effect of connectivity treatment on ZB, but αS, αS_PIE_ and J were not significantly affected. Consequently, we did not build a complex SEM afterward. The results of each initial SEM are presented in the supplementary material (Figure S2-S3, Table S11-S12). SEMs were implemented using the ‘semEff’ v. 0.6.1 package (Murphy, 2022).

The effect of initial mean regional abundance (i.e., mean abundance of the given ASV within a metacommunity in week 4, RA), ZB in week 16, and the fragmentation treatment on the survival probability of the individual prokaryote and microeukaryote ASVs over the entire experimental period (i.e., presence or absence of the given ASV in week 16) was studied with generalized linear mixed-effects models (GLMMs). These GLMMs were built with binomial function, where RA, ZB and treatment were included as fixed, while metacommunity ID as a nested random effect factor. Both for prokaryotes and microeukaryotes, we compared a model without and including the interaction between RA and treatment as well as between ZB and treatment with chi-square test using the anova() function. As interaction terms significantly improved the model fits (prokaryotes: χ^2^(2, 31808) = 25.099; P<0.001; microeukaryotes: χ^2^(2, 35550) = 7.8203; P<0.05), we retained both of them in both models. RA and ZB were z-score standardized to eliminate the bias that can arise from the different measurement units and scales. Odds ratios and 95% confidence intervals were calculated using the tab_model() function of ‘sjPlot’ v. 2.8.15 package (Lüdecke, 2023).

In the main part of the paper, we present the results obtained for prokaryotes and microeukaryotes at the last sampling event. All other results were provided in the supplementary material. Although whisker plots and scatter plots were created based on raw datasets for an easier visual comparison of different organism groups and sampling events, when necessary, dependent variables were transformed to achieve normal distribution. Applied transformations are presented in the relevant tables along with the results of statistical tests (Table S3, S6, S11-S18).

## Results

Local richness of microeukaryotes, i.e., both αS and αS_PIE_ were significantly lower in the fragmented metacommunities compared to the connected ones at the end of the experiment (week 16; Figure 2, S4-S5, Table S13). In prokaryotes, a significant negative effect of fragmentation on αS emerged by week 12, while it disappeared by week 16 with no significant effect on αS and αS_PIE_. Fragmentation resulted in a significantly lower J in microeukaryotes by the end of the experiment (Figure 2, S6, Table S13). In contrast, prokaryote J was not affected by treatment. Similarly, we found a negative effect of fragmentation on microeukaryote αS but no effect on prokaryotes when analyzing the entire experimental period (Table S14).

**Figure 2.**
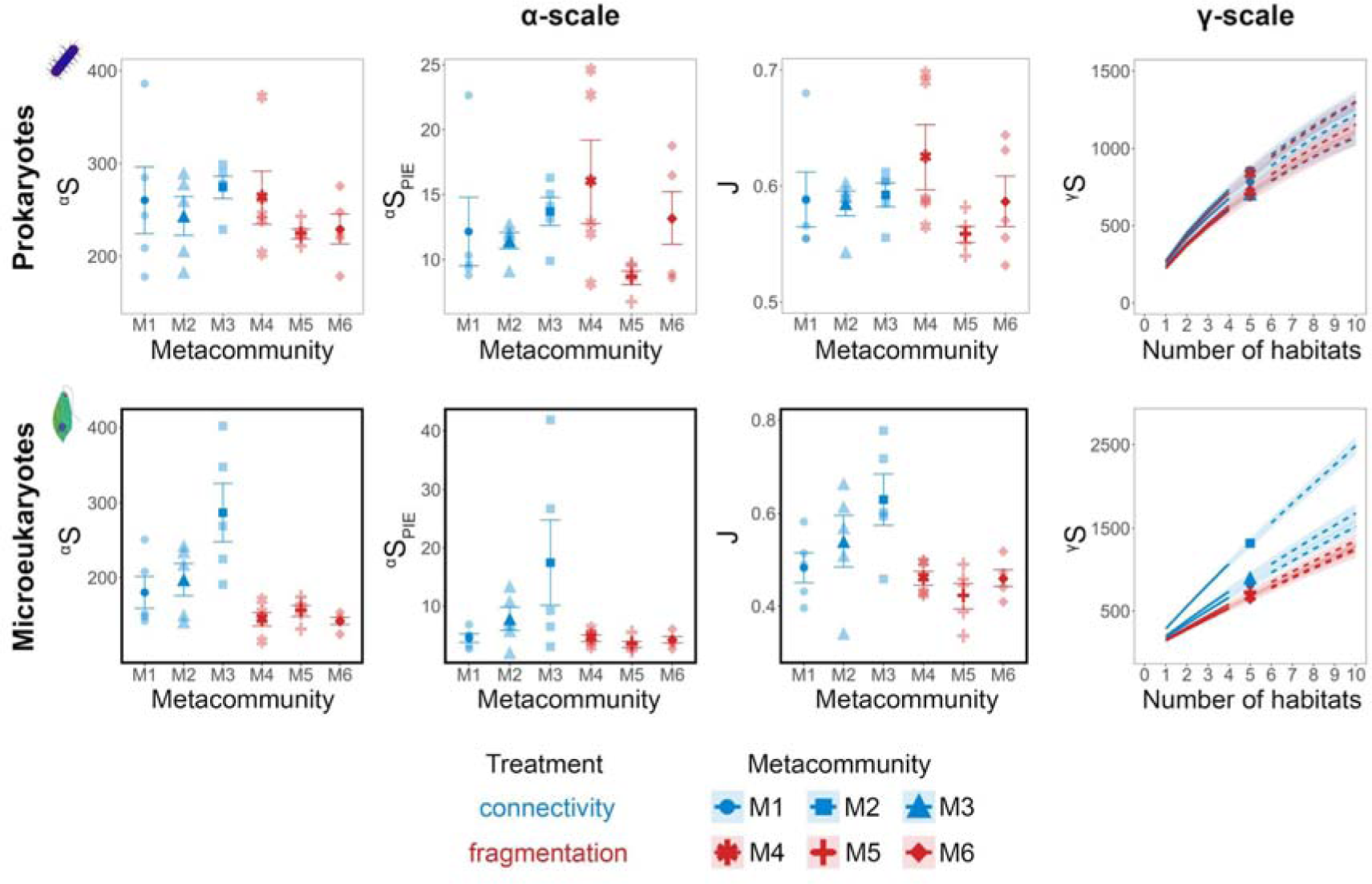
Number of observed ASVs (αS), effective number of ASVs of PIE (αS_PIE_) and evenness (J) at α-scale, and accumulation curves with the number of observed ASVs (γS) at γ-scale for prokaryotes and microeukaryotes in each metacommunity at the end of the experiment (week 16). Along with the accumulation curves, extrapolated estimates (dashed lines) and 95% confidence intervals (error bands) are also displayed. Boxes marked with a bold black frame indicate a significant treatment effect (P<0.05) resulting from the ANOVA.

We did not find a significant treatment effect on β_S_ in microeukaryotes (Figure S7, Table S13). In prokaryotes, β_S_ was significantly lower in the connected metacommunities in week 12, but even this difference disappeared by the end of the experiment (Figure S7, Table S13).

At the regional scale (γ), fragmentation resulted in a lower number of ASVs in microeukaryotes by the end of the experiment. This was evidenced by the accumulation curves and no overlap between the 95% confidence intervals between treatments (Figure 2, Figure S8). In prokaryotes, no treatment effect on γS was found for the entire experimental duration (Figure 2, Figure S8).

Chl-a was not affected significantly by the treatment (Table S15). αS_PIE_ of phytoplankton (based on microscopic identification) was significantly decreased by the fragmentation treatment at the end of the experiment, while we found no effect on αS and J (Table S16), similarly to the results obtained for the microeukaryote communities based on metabarcoding data.

Prokaryote αS, αS_PIE_ and J were not affected significantly by either treatment or ZB at the end of the experiment (week 16) based on the LMM and SEM (Figure 3, S2, Table S11, S17). In microeukaryotes, αS had a positive relationship with ZB in the connected metacommunities and a negative relationship in the fragmented ones (Figure 3, Table S17; ANOVA: P_Treatment*log(ZB)_<0.05, F_Treatment*log(ZB)_=7.144). Microeukaryote αS_PIE_ and J showed an increase with the increase of ZB irrespective of treatment (Figure 3, Table S17).

**Figure 3.**
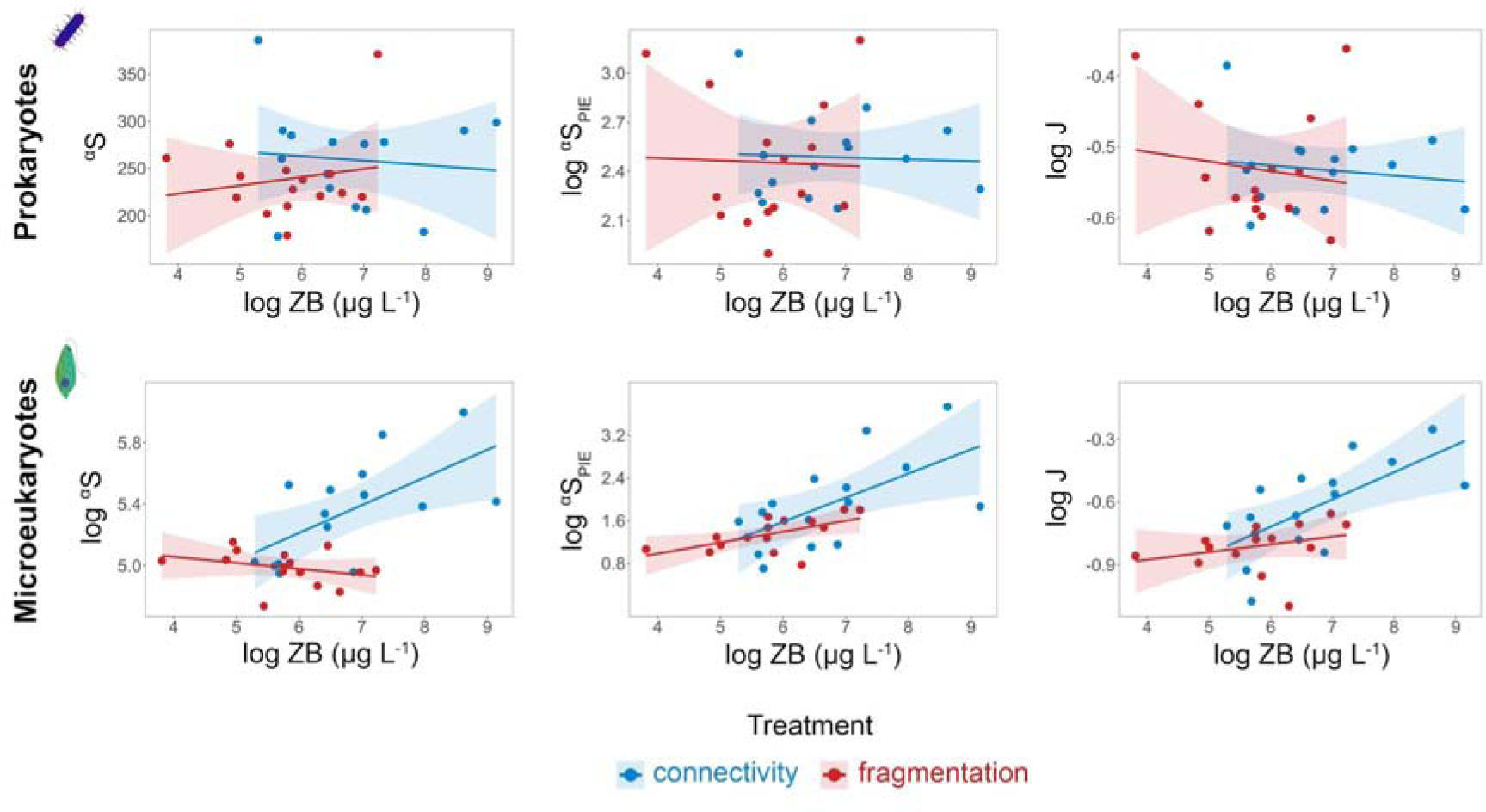
Linear regression plots illustrating the change in the number of observed ASVs (αS), effective number of ASVs of PIE (αS_PIE_), and evenness (J) as a function of zooplankton biomass (ZB) in the two treatment groups (connectivity and fragmentation) for prokaryotes and microeukaryotes at the end of the experiment (week 16). Solid lines represent the fitted linear models (error bands: standard errors). ZB, αS_PIE_ and J were log-transformed in both cases, and αS in the case of microeukaryotes. Summary statistics of the LMMs, including metacommunity ID as a random factor are presented in the supplementary material (Table S17).

Based on the SEM (Figure 4, Table S18), fragmentation had a significant negative direct effect on microeukaryote αS, while the direct effect of microeukaryote αS on ZB, and of ZB on microeukaryote J was a significant positive effect. Although treatment directly did not increase ZB significantly, its indirect negative effect including microeukaryote αS as a mediator was significant, which is in line with the negative relationship between microeukaryote αS and ZB under fragmentation (Figure 3). Microeukaryote J significantly decreased as a response to fragmentation and in this case, the direct treatment effect was stronger than the indirect (i.e., the effect of treatment on microeukaryote J with the mediation of microeukaryote αS and ZB) (Figure 4, Table S18).

**Figure 4.**
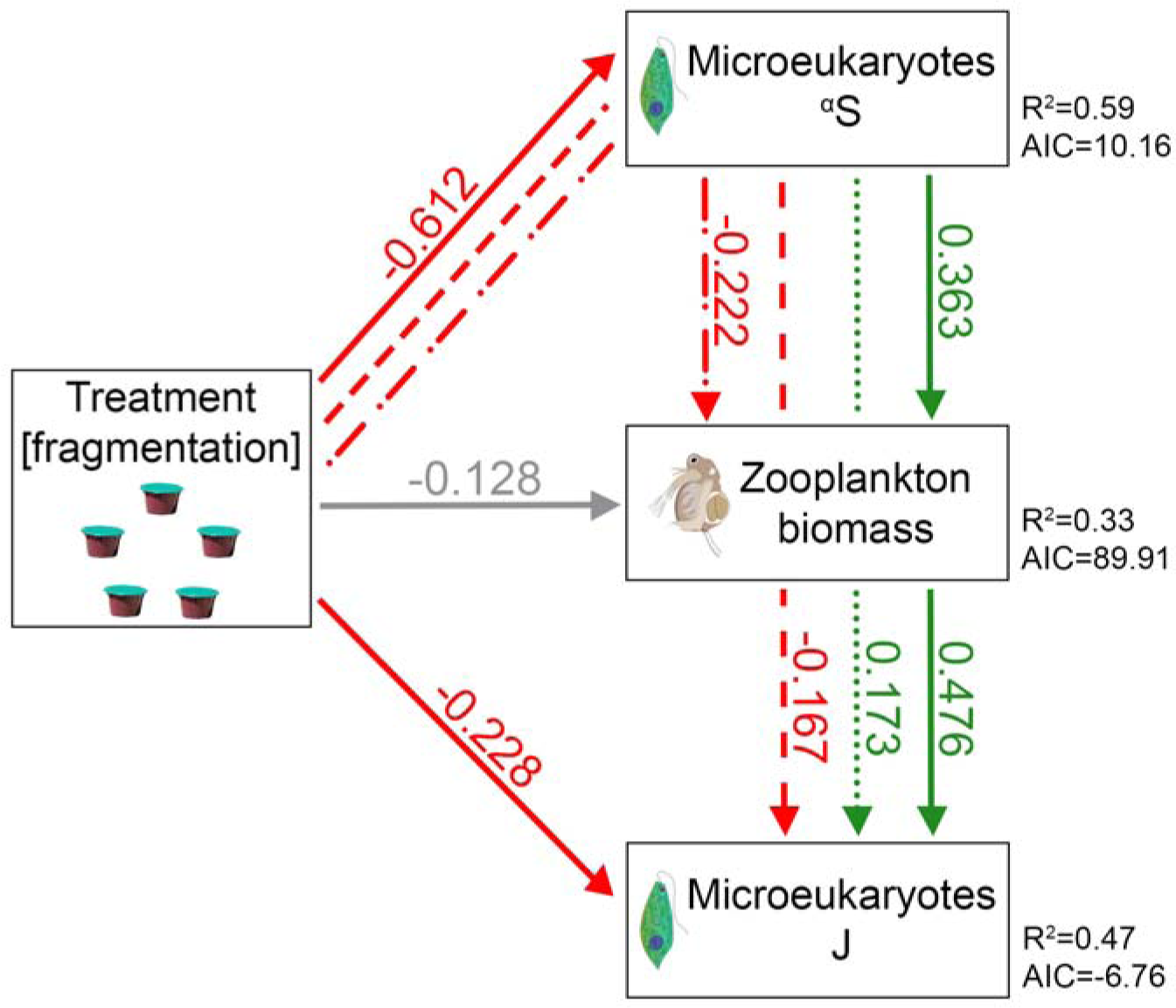
Structural equation model (SEM) showing the relationships (direct and indirect effects) between treatment, zooplankton biomass and diversity (αS or J) in microeukaryotes at the end of the experiment (week 16). Variables are represented by boxes and the directional relationships by single-head arrows (solid line: direct effect; dashed, dotted and dash-dotted lines: indirect effects; green: significant positive effect; red: significant negative effect; gray: non-significant effect). Standardized path coefficients are shown on the arrows. R^2^ and AIC values for the component models are indicated next to the boxes of endogenous variables. αS, zooplankton biomass and J were log-transformed prior to the analysis. In the models, microeukaryote αS, zooplankton biomass and treatment were included as fixed, and metacommunity ID as a random effect factor nested within treatment.

The initial mean regional abundance significantly affected the survival probability of both prokaryotes and microeukaryotes (based on GLMM with an odds ratio>1). Besides the regional abundance, ZB in week 16, treatment and also their interaction significantly affected the survival probability of microeukaryote ASVs. These results indicate that ZB had different effects in the two treatments: an increase in ZB increased the survival probability in microeukaryotes in the connected metacommunities while decreasing it in the fragmented ones (Figure 5, Table S19). In prokaryotes, the treatment did not affect the survival probability, however, the interaction effect of regional abundance and treatment was significant. This indicates that an increase in the regional abundance increased the survival probability in the connected metacommunities to a lesser extent than in the fragmented metacommunities (Figure 5, Table S19).

**Figure 5.**
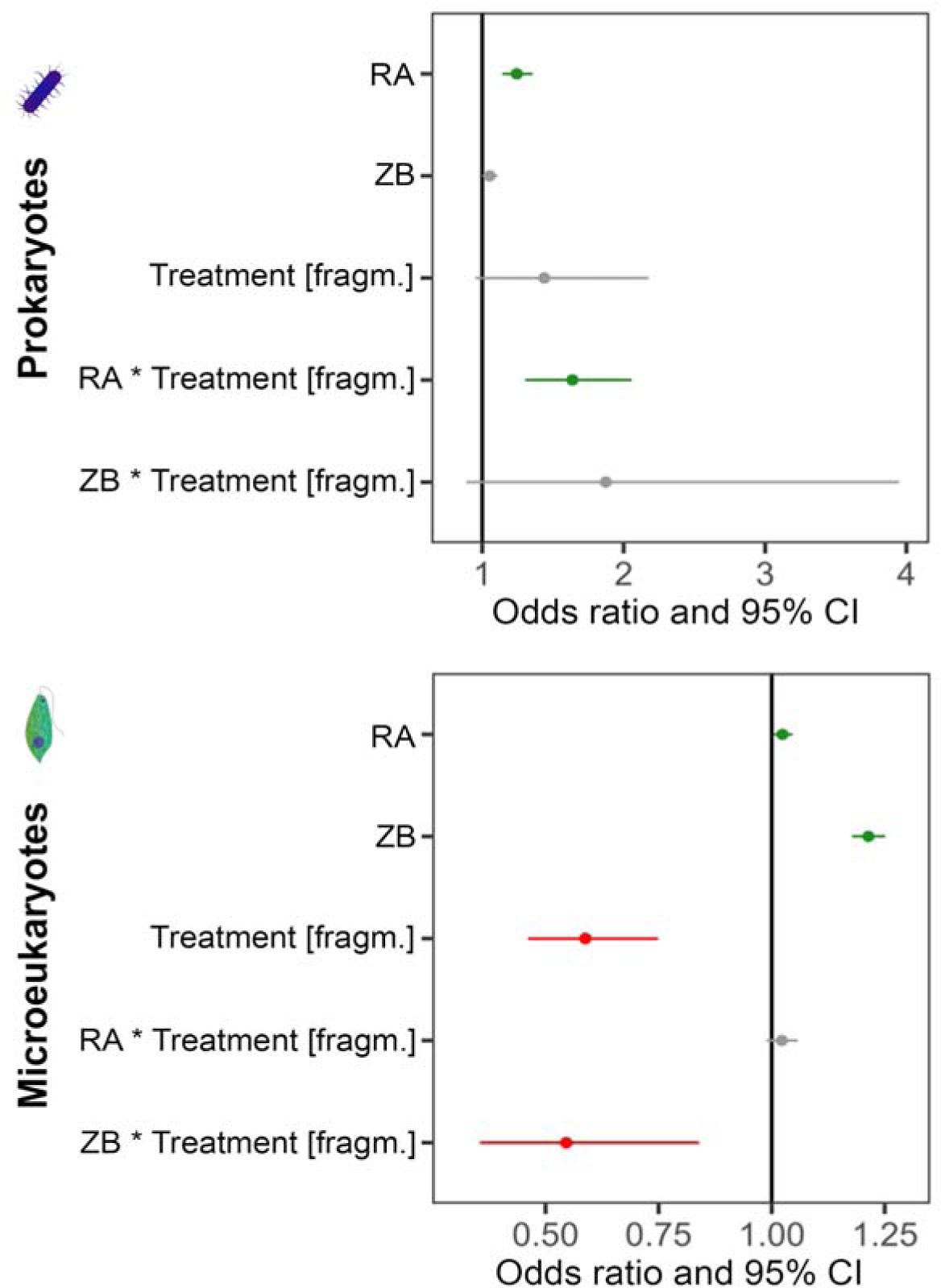
Forest plots demonstrating the effect of the initial mean regional abundance (RA), zooplankton biomass (ZB) in week 16, treatment, and the interaction of treatment with RA and ZB on the survival probability of individual ASVs by the end of the experiment (red: significant negative, green: significant positive effect; gray: non-significant effect). The plots are based on odds ratios (closed circles, with whiskers representing 95% confidence intervals) in the generalized linear mixed-effects models (GLMMs) performed on the presence/absence of the ASVs in week 16 in prokaryotes and microeukaryotes. In the models, RA, ZB and treatment were included as fixed, and metacommunity ID as a random effect factor nested within treatment.

## Discussion

We provided clear experimental evidence on the negative effects of habitat fragmentation, specifically its connectivity loss component, on local and regional biodiversity. In our study based on experimental pond networks, we found that without changes in underlying local environmental conditions or loss of total habitat amount, connectivity loss alone exerted a strong negative effect on both local and regional richness and evenness of unicellular microeukaryotes. This negative effect could be observed both in αS and αS_PIE_, indicating that rare and abundant taxa were all affected. In line with our expectation, diversity of prokaryotes was less affected, although the initial mean regional abundance of prokaryote taxa significantly affected their survival probability indicating a potential extinction debt. We also found that the negative effect of connectivity loss was partly related to the dispersal-mediated trophic interactions with grazers. Overall, our study clearly demonstrates that loss of connectivity within a pond network can lead to a significant loss of microbial biodiversity.

Given the absence of environmental gradients, our experimental metacommunity can be considered as representing a case where patch dynamics is the major process of community assembly. In such metacommunities, interspecific interactions play a predominant role in community assembly modulated by dispersal as a key process (Leibold et al., 2004; Thompson et al., 2020). In our study, we expected a direct effect of habitat fragmentation in the form of connectivity loss especially in the microeukaryote metacommunities. Indeed, by the end of the 16-week experiment, we found significantly lower local richness of microeukaryotes in the fragmented metacommunities compared to the connected ones. This was apparent both in αS and αS_PIE_, indicating a negative effect on both rare and abundant taxa. A similar difference also emerged at the regional scale, resulting in lower regional diversity. Overall, these results clearly point out that the disruption of weekly dispersal was able to lead to the loss of several taxa. These findings were further supported by our analysis on direct and indirect effects and also by a separate analysis based on survival probabilities, providing further evidence for the negative effect of connectivity loss on the survival of microeukaryotes during the course of the experiment.

Besides local richness, evenness of the microeukaryote communities under connectivity loss was also significantly lower compared to those where connectivity was maintained until the end of the experiment. Our results support that connectivity can alter interspecific interactions by facilitating immigration of species from neighboring habitats, thereby decreasing the local dominance of competitively superior species (Leibold et al., 2004). Habitats affected by connectivity loss may exhibit reduced evenness as competitively inferior species may get outcompeted due to the reduced efficiency of rescue effects via immigration (Brown and Kodric-Brown, 1977; Dey and Joshi, 2013; Eriksson et al., 2014; Marini et al., 2014). Furthermore, local survival probabilities can strongly depend on initial differences in population size (Kim et al., 2016; Matthies et al., 2004), as it was also supported by our analyses. Given that initial mean regional abundance was an important predictor of survival of microeukaryotes in both treatments (connectivity vs. fragmentation), this might also point at the role of priority effects to some extent. These strong dominance patterns were enhanced by both the direct and indirect effects (by decreasing the biomass of zooplankton grazers via reducing microeukaryote richness) of connectivity loss and in the fragmentation treatment.

Besides its effect on the local scale, habitat fragmentation also led to significantly lower regional richness compared to those metacommunities where continuous connectivity ensured the dispersal of organisms evidenced by the species accumulation curves. This was in agreement with our prediction based on earlier theoretical (Thompson et al., 2017) and experimental studies (Gilbert et al., 1998). The effects of connectivity loss on regional biodiversity can be less straightforward in more complex natural settings, involving heterogeneous habitat patches and complex food webs (e.g., Evans et al., 2017; Gibb and Hochuli, 2002; Gibson et al., 2013). Our results, however, suggested the loss in regional diversity under weakened rescue effects due to the disruption of the dispersal network.

While homogeneity of the abiotic environment and phytoplankton biomass (approximated by Chl-a fluorescence) persisted during the experiment, indicated by no significant differences among treatments, fragmentation reduced ZB by the end of the experiment. This effect on ZB was not coupled with a parallel effect on zooplankton taxon richness (Table S6) and it indicates potential differences in the biotic interactions between the treatments. A general negative effect on both the diversity of microeukaryotes and the biomass of their zooplankton grazers may be a result (and actually a combination) of three processes. First, connectivity loss might disadvantage both microeukaryotes and zooplankton directly. A direct effect seems to be the case for microeukaryotes, supported by a series of complementary analyses of both local and regional diversity, including the survival probability analysis showing both the importance of initial mean regional abundances and the effect of connectivity loss. The negative effect of fragmentation on ZB, at the same time, was an indirect effect (Figure 4). The lack of direct significant effect on zooplankton taxon richness could have been also linked to differences in taxonomic resolution used for the two groups (metabarcoding: total of 11,134 ASVs for microeukaryotes, taxon richness via microscopic analysis for zooplankton: 16 taxa). However, the standard microscopic analysis of phytoplankton (which was more comparable to the microscopic analysis of zooplankton in terms of taxonomic resolution, with a total of 82 taxa encountered during the experiment) largely corroborated the metabarcoding-based results and led us to the same conclusion. These results therefore rather indicate potentially different trophic relationships under fragmentation.

Second, phytoplankton biomass (Chl-a fluorescence; Table S15) did not differ across the treatments, hence the quantity of algal food for zooplankton was presumably similar in both treatments. In contrast to the fragmented metacommunities, maintaining connectivity in the connectivity treatment could have indirectly contributed to higher ZB via an increased diversity of their food resource (Striebel et al., 2012, Marzetz et al., 2017). This was indeed supported by the results of our complex structural equation model, where fragmentation had both a significant direct effect on microeukaryote αS and no direct, but an indirect effect on ZB via mediation of local diversity in microeukaryotes.

Third and finally, changes in the biotic interactions between zooplankton grazers and microeukaryotes might have also contributed to the changes in local diversity patterns via another indirect pathway. Our results suggest that the lower evenness of microeukaryotes under fragmentation resulted both from the direct negative effect of the treatment and the effect of a lower ZB. In the connectivity treatment, the higher biomass of the relatively small-sized cladocerans (*Moina* and *Macrothrix*), being dominant in the community, seemingly exerted an overall positive grazing effect on microeukaryotes. Intermediate levels of grazing can be considered as a form of intermediate disturbance (Colburn, 2008; Joubert et al., 2017; Shea et al., 2004; Yuan et al., 2016). It is possible that the dominant cladocerans exerted such an effect, which was supported by the positive relationship of ZB with both microeukaryote diversity and evenness in the connectivity treatment (Figure 3). Zooplankton communities dominated by large-bodied generalist cladocerans such as *Daphnia magna* might exert different effects on their prey communities, e.g., even masking the positive effect of connectivity on local diversity (Berga et al., 2015). In contrast, the small-sized cladocerans and rotifers in our experiment likely contributed to decreasing the dominance of a selected set of otherwise superior competitor species either via more selective grazing (based on size, taste, concentration, nutritional value, etc.; Sterner 1989) or by a general but intermediate grazing pressure that could benefit the competitively inferior species (Lubchenco, 1978; Pulungan et al., 2019). The pattern we found that evenness might be linked to treatment-specific differences in grazing effects was also confirmed by our microscopic analyses that showed a higher dominance of picophytoplankton in the fragmentation treatment (Table S20, Figure S9). These small sized unicellular algae are within the filtration range of *Moina* (2-40 µm; Pagano, 2008) and several rotifers (Pourriot, 1977; Rothhaupt, 1990), thus, the high biomass of these zooplankton taxa at the end of the experiment seemed to be linked to the decline of pico-size algae in the connected mesocosms.

Both pro-and microeukaryotes are known to be efficient passive dispersers across long distances (Genitsaris et al., 2011; Luef et al., 2007; Mony et al., 2020; Szabó et al., 2022) and their communities often show no or only a weak spatial structuring (Barta et al., 2024; Beisner et al., 2006; Padial et al., 2014). At the same time, there is also increasing evidence for the importance of connectivity for sustaining biodiversity and temporal stability of local communities in these groups (e.g., Berga et al., 2015; Engel et al., 2020; Guelzow et al., 2017; Thompson et al., 2015; Vad et al., 2023). Among the two, prokaryotes have been considered to be less dispersal limited due to their smaller size (De Bie et al., 2012), their larger population sizes of a few orders of magnitude compared to microeukaryotes (Berninger et al., 1991; De Bie et al., 2012; Fenchel, 1988), and their higher rate of propagule production (i.e. shorter generation time) (Zubkov, 2014). In line with these, we predicted a comparably smaller effect of fragmentation on prokaryotes than on microeukaryotes. This was supported by our results based on the analyses of both local and regional diversity and survival probabilities, with no significant direct effect of fragmentation on prokaryotes. On the other hand, mean regional abundance at the start of the fragmentation treatment was an even stronger predictor of local survival in prokaryotes than in microeukaryotes, especially when predicting survival in the fragmentation treatment. This might point at higher levels of monopolization in the absence of dispersal (Loeuille and Leibold, 2008; Urban and De Meester, 2009) and a potentially longer time frame that would have been necessary to detect changes in species richness. Larger populations or regionally more frequent species of a metacommunity are less prone to extinction driven by stochastic events (De Silva and Leimgruber, 2019; Eriksson et al., 2014; Kim et al., 2016; Lande et al., 2003; Matthies et al., 2004). ASVs with lower initial mean abundance might also experience slow declines over a longer time frame, eventually leading to extinction events also in prokaryotes. This delayed response might indicate a potential extinction debt in this group as a response to fragmentation.

An important novelty of our study lies in the way of application of the fragmentation treatment, which was independent of habitat amount reduction: we modeled connected sites that later got disconnected, thereby mimicking the natural dynamics of connectivity loss in a landscape. Furthermore, in contrast to several previous studies that applied a constant dispersal from an external regional species pool, we maintained a within-network connectivity, thus avoiding bias due to the potential mass effects and a potential loss in species due to a “mesocosm effect”; this occurs when species are lost during the first phase of an experiment due to the recent inoculation from natural ponds, which somewhat differ in their local characteristics from the experimental conditions (see e.g., Adey et al., 1996; Limberger et al., 2019; Williams, 2002). Such effects would have made it more difficult to track treatment-specific extinction events. Nevertheless, the results we found also indicate that longer time (even months) might be needed for a detectable signal of fragmentation in such an experimental setup.

Habitat fragmentation can negatively affect biodiversity and alter ecosystem functioning in several direct and indirect ways. Although relatively much is known about its effects on terrestrial (Gonzalez et al., 1998; Graham et al., 2022; Haddad et al., 2017) and aquatic macroorganisms (Hamer, 2016; Hamer et al., 2023; Johnson et al., 2013; Rösch et al., 2013), aquatic prokaryotes and microeukaryotes, most of which act like important producers or decomposers in the food web, are still understudied in this regard. Our experimental study demonstrates that connectivity loss, i.e., disruption of dispersal among habitat patches plays a crucial role by directly reducing both local and regional richness as well as community evenness. Our results were especially explicit in unicellular microeukaryotes, which may be linked to their smaller population sizes (Berninger et al., 1991; De Bie et al., 2012; Fenchel, 1988) and longer generation times (Zubkov, 2014) compared to prokaryotes. We also found strong evidence that connectivity loss can indirectly decrease microeukaryote evenness via modulating biotic interactions, more precisely, the strength of zooplankton grazing effect. Overall, these results highlight that maintaining connectivity in networks of such highly sensitive small waterbodies as ponds is essential to preserve microbial diversity and ecosystem functioning. Furthermore, we emphasize that for better understanding the direct and indirect processes resulting from connectivity loss, and thus being able to assist conservation strategies and management practices, longer-term experiments incorporating complex food webs are required.

## Acknowledgements

We acknowledge the practical help of Anna Dervalics, Csenge Póda, Dóra Fehér, Ivana Lukić, Kamilla Nyitrai, Kinga Glatz, Máté Burányi, Olivér Payer, Zorka Szabadkai, Virág Csiszár, and Vivien Kardos who participated in running the experiment and collecting the samples. We especially thank Bernadett Szabó and Csilla Laskai for their help in the laboratory processing of microbial samples, Attila Szabó for his guidance in amplicon data processing, and Katalin Bodolai for measuring the chemical parameters of water samples. We are grateful to Veronika Bókony whose suggestions helped improve the statistical methods.

## Data availability statement

The data that support the findings of this study are openly available in the European Nucleotide Archive at https://www.ebi.ac.uk/ena/browser/view/PRJEB78363, reference number PRJEB78363.

## References

Adey, W.H., Finn, M., Kangas, P., Lange, L., Luckett, C., Spoon, D.M., 1996. A Florida Everglades Mesocosm — model veracity after four years of self-organization. Ecol. Eng. 6, 171–224. 10.1016/0925-8574(95)00057-7

Baas Becking, L.G.M., Nicolai, E., 1934. On the ecology of a Sphagnum Bog. Blumea: Biodiversity, Evolution and Biogeography of Plants 1, 10–45.

Barta, B., Szabó, A., Szabó, B., Ptacnik, R., Vad, C.F., Horváth, Z., 2024. How pondscapes function: connectivity matters for biodiversity even across small spatial scales in aquatic metacommunities. Ecography 2024, e06960. 10.1111/ecog.06960

Bates, D., Mächler, M., Bolker, B., Walker, S., 2015. Fitting Linear Mixed-Effects Models Using **lme4**. J. Stat. Softw. 67. 10.18637/jss.v067.i01

Beisner, B.E., Peres-Neto, P.R., Lindström, E.S., Barnett, A., Longhi, M.L., 2006. THE ROLE OF ENVIRONMENTAL AND SPATIAL PROCESSES IN STRUCTURING LAKE COMMUNITIES FROM BACTERIA TO FISH. Ecology 87, 2985–2991. 10.1890/0012-9658(2006)87[2985:TROEAS]2.0.CO;2

Berga, M., Östman, Ö., Lindström, E.S., Langenheder, S., 2015. Combined effects of zooplankton grazing and dispersal on the diversity and assembly mechanisms of bacterial metacommunities. Environ. Microbiol. 17, 2275–2287. 10.1111/1462-2920.12688

Berninger, U., Finlay, B.J., KuuppoCLeinikki, P., 1991. Protozoan control of bacterial abundances in freshwater. Limnol. Oceanogr. 36, 139–147. 10.4319/lo.1991.36.1.0139

Boros, E., V.-Balogh, K., Vörös, L., Horváth, Z., 2017. Multiple extreme environmental conditions of intermittent soda pans in the Carpathian Basin (Central Europe). Limnologica 62, 38–46. 10.1016/j.limno.2016.10.003

Briggs, C.J., Hoopes, M.F., 2004. Stabilizing effects in spatial parasitoid–host and predator– prey models: a review. Theor. Popul. Biol. 65, 299–315. 10.1016/j.tpb.2003.11.001

Brooks, T.M., Mittermeier, R.A., Mittermeier, C.G., Da Fonseca, G.A.B., Rylands, A.B., Konstant, W.R., Flick, P., Pilgrim, J., Oldfield, S., Magin, G., HiltonCTaylor, C., 2002. Habitat Loss and Extinction in the Hotspots of Biodiversity. Conserv. Biol. 16, 909–923. 10.1046/j.1523-1739.2002.00530.x

Brown, J.H., Kodric-Brown, A., 1977. Turnover Rates in Insular Biogeography: Effect of Immigration on Extinction. Ecology 58, 445–449. 10.2307/1935620

Chao, A., Gotelli, N.J., Hsieh, T.C., Sander, E.L., Ma, K.H., Colwell, R.K., Ellison, A.M., 2014. Rarefaction and extrapolation with Hill numbers: a framework for sampling and estimation in species diversity studies. Ecol. Monogr. 84, 45–67. 10.1890/13-0133.1

Chase, J.M., 2003. Community assembly: when should history matter? Oecologia 136, 489–498. 10.1007/s00442-003-1311-7

Chase, J.M., Jeliazkov, A., Ladouceur, E., Viana, D.S., 2020. Biodiversity conservation through the lens of metacommunity ecology. Ann. N. Y. Acad. Sci. 1469, 86–104. 10.1111/nyas.14378

Colburn, E.A., 2008. Temporary Waters, in: Encyclopedia of Ecology. Elsevier, pp. 657–670. 10.1016/B978-0-444-63768-0.00361-9

De Bie, T., De Meester, L., Brendonck, L., Martens, K., Goddeeris, B., Ercken, D., Hampel, H., Denys, L., Vanhecke, L., Van der Gucht, K., Van Wichelen, J., Vyverman, W., Declerck, S.A.J., 2012. Body size and dispersal mode as key traits determining metacommunity structure of aquatic organisms. Ecol. Lett. 15, 740–747. 10.1111/j.1461-0248.2012.01794.x

de Boer, M.K., Moor, H., Matthiessen, B., Hillebrand, H., Eriksson, B.K., 2014. Dispersal restricts local biomass but promotes the recovery of metacommunities after temperature stress. Oikos 123, 762–768. 10.1111/j.1600-0706.2013.00927.x

De Silva, S., Leimgruber, P., 2019. Demographic Tipping Points as Early Indicators of Vulnerability for Slow-Breeding Megafaunal Populations. Front. Ecol. Evol. 7, 171. 10.3389/fevo.2019.00171

Dey, S., Joshi, A., 2013. Effects of constant immigration on the dynamics and persistence of stable and unstable Drosophila populations. Sci. Rep. 3, 1405. 10.1038/srep01405

Eggers, S.L., Eriksson, B.K., Matthiessen, B., 2012. A heat wave and dispersal cause dominance shift and decrease biomass in experimental metacommunities. Oikos 121, 721–733. 10.1111/j.1600-0706.2011.19714.x

Engel, F.G., Matthiessen, B., Eriksson, B.K., 2020. A heatwave increases turnover and regional dominance in microbenthic metacommunities. Basic Appl. Ecol. 47, 1–11. 10.1016/j.baae.2020.03.003

Eriksson, A., Elías-Wolff, F., Mehlig, B., Manica, A., 2014. The emergence of the rescue effect from explicit within- and between-patch dynamics in a metapopulation. Proc. R. Soc. B Biol. Sci. 281, 20133127. 10.1098/rspb.2013.3127

Evans, M.J., Banks, S.C., Driscoll, D.A., Hicks, A.J., Melbourne, B.A., Davies, K.F., 2017. ShortC and longCterm effects of habitat fragmentation differ but are predicted by response to the matrix. Ecology 98, 807–819. 10.1002/ecy.1704

Fahrig, L., 2019. Habitat fragmentation: A long and tangled tale. Glob. Ecol. Biogeogr. 28, 33–41. 10.1111/geb.12839

Fahrig, L., 2017. Ecological Responses to Habitat Fragmentation Per Se. Annu. Rev. Ecol. Evol. Syst. 48, 1–23. 10.1146/annurev-ecolsys-110316-022612

Fahrig, L., 2013. Rethinking patch size and isolation effects: the habitat amount hypothesis. J. Biogeogr. 40, 1649–1663. 10.1111/jbi.12130

Fahrig, L., 2002. EFFECT OF HABITAT FRAGMENTATION ON THE EXTINCTION THRESHOLD: A SYNTHESIS *. Ecol. Appl. 12, 346–353. 10.1890/1051-0761(2002)012[0346:EOHFOT]2.0.CO;2

Fenchel, T., 1988. MARINE PLANKTON FOOD CHAINS. Annu. Rev. Ecol. Syst. 19, 19–38. 10.1146/annurev.es.19.110188.000315

Finlay, B.J., 2002. Global Dispersal of Free-Living Microbial Eukaryote Species. Science 296, 1061–1063. 10.1126/science.1070710

Fletcher, R.J., Didham, R.K., Banks-Leite, C., Barlow, J., Ewers, R.M., Rosindell, J., Holt, R.D., Gonzalez, A., Pardini, R., Damschen, E.I., Melo, F.P.L., Ries, L., Prevedello, J.A., Tscharntke, T., Laurance, W.F., Lovejoy, T., Haddad, N.M., 2018. Is habitat fragmentation good for biodiversity? Biol. Conserv. 226, 9–15. 10.1016/j.biocon.2018.07.022

Foissner, W., 2006. Biogeography and dispersal of micro-organisms: a review emphasizing prostists. Acta Protozool. 45, 111–136.

Frankham, R., 2006. Genetics and landscape connectivity. In K. R. Crooks, & M. Sanjayan (Eds.), Connectivity conservation (pp. 72-96). (Conservation biology series; Vol. 14). Cambridge University Press (CUP). 10.2277/0521857066

Genitsaris, S., Moustaka-Gouni, M., Kormas, K., 2011. Airborne microeukaryote colonists in experimental water containers: diversity, succession, life histories and established food webs. Aquat. Microb. Ecol. 62, 139–152. 10.3354/ame01463

Gianuca, A.T., Declerck, S.A.J., Lemmens, P., De Meester, L., 2017. Effects of dispersal and environmental heterogeneity on the replacement and nestedness components of βCdiversity. Ecology 98, 525–533. 10.1002/ecy.1666

Gibb, H., Hochuli, D.F., 2002. Habitat fragmentation in an urban environment: large and small fragments support different arthropod assemblages. Biol. Conserv. 106, 91–100. 10.1016/S0006-3207(01)00232-4

Gibson, L., Lynam, A.J., Bradshaw, C.J.A., He, F., Bickford, D.P., Woodruff, D.S., Bumrungsri, S., Laurance, W.F., 2013. Near-Complete Extinction of Native Small Mammal Fauna 25 Years After Forest Fragmentation. Science 341, 1508–1510. 10.1126/science.1240495

Gilbert, F., Gonzalez, A., Evans-Freke, I., 1998. Corridors maintain species richness in the fragmented landscapes of a microecosystem. Proc. R. Soc. Lond. B Biol. Sci. 265, 577–582. 10.1098/rspb.1998.0333

Gonzalez, A., Lawton, J.H., Gilbert, F.S., Blackburn, T.M., Evans-Freke, I., 1998. Metapopulation Dynamics, Abundance, and Distribution in a Microecosystem. Science 281, 2045–2047. 10.1126/science.281.5385.2045

Gonzalez, A., Rayfield, B., Lindo, Z., 2011. The disentangled bank: How loss of habitat fragments and disassembles ecological networks. Am. J. Bot. 98, 503–516. 10.3732/ajb.1000424

Graham, C.D.K., Warneke, C.R., Weber, M., Brudvig, L.A., 2022. The impact of habitat fragmentation on domatia-dwelling mites and a mite-plant-fungus tritrophic interaction. Landsc. Ecol. 37, 3029–3041. 10.1007/s10980-022-01529-2

Guelzow, N., Muijsers, F., Ptacnik, R., Hillebrand, H., 2017. Functional and structural stability are linked in phytoplankton metacommunities of different connectivity. Ecography 40, 719–732. 10.1111/ecog.02458

Haddad, N.M., Brudvig, L.A., Clobert, J., Davies, K.F., Gonzalez, A., Holt, R.D., Lovejoy, T.E., Sexton, J.O., Austin, M.P., Collins, C.D., Cook, W.M., Damschen, E.I., Ewers, R.M., Foster, B.L., Jenkins, C.N., King, A.J., Laurance, W.F., Levey, D.J., Margules, C.R., Melbourne, B.A., Nicholls, A.O., Orrock, J.L., Song, D.-X., Townshend, J.R., 2015. Habitat fragmentation and its lasting impact on Earth’s ecosystems. Sci. Adv. 1, e1500052. 10.1126/sciadv.1500052

Haddad, N.M., Gonzalez, A., Brudvig, L.A., Burt, M.A., Levey, D.J., Damschen, E.I., 2017. Experimental evidence does not support the Habitat Amount Hypothesis. Ecography 40, 48–55. 10.1111/ecog.02535

Hagen, M., Kissling, W.D., Rasmussen, C., De Aguiar, M.A.M., Brown, L.E., Carstensen, D.W., Alves-Dos-Santos, I., Dupont, Y.L., Edwards, F.K., Genini, J., Guimarães, P.R., Jenkins, G.B., Jordano, P., Kaiser-Bunbury, C.N., Ledger, M.E., Maia, K.P., Marquitti, F.M.D., Mclaughlin, Ó., Morellato, L.P.C., O’Gorman, E.J., Trøjelsgaard, K., Tylianakis, J.M., Vidal, M.M., Woodward, G., Olesen, J.M., 2012. Biodiversity, Species Interactions and Ecological Networks in a Fragmented World, in: Advances in Ecological Research. Elsevier, pp. 89–210. 10.1016/B978-0-12-396992-7.00002-2

Hamer, A.J., 2016. Accessible habitat delineated by a highway predicts landscape-scale effects of habitat loss in an amphibian community. Landsc. Ecol. 31, 2259–2274. 10.1007/s10980-016-0398-2

Hamer, A.J., Mechura, T., Puky, M., 2023. Patterns in usage of under-road tunnels by an amphibian community highlights the importance of tunnel placement and design for mitigation. Glob. Ecol. Conserv. 43, e02420. 10.1016/j.gecco.2023.e02420

Hanski, I., 2011. Habitat Loss, the Dynamics of Biodiversity, and a Perspective on Conservation. AMBIO 40, 248–255. 10.1007/s13280-011-0147-3

Harrisson, K.A., Pavlova, A., Amos, J.N., Takeuchi, N., Lill, A., Radford, J.Q., Sunnucks, P., 2012. Fine-scale effects of habitat loss and fragmentation despite large-scale gene flow for some regionally declining woodland bird species. Landsc. Ecol. 27, 813–827. 10.1007/s10980-012-9743-2

Hassell, M.P., Godfray, H.C., Comins, H.N., 1993. Effects of global change on the dynamics of insect host-parasitoid interactions. – In: Kareiva, P.M., Kingsolver, J.G., Huey, R.B. (editors), Biotic interactions and global change. Sinauer Associates Inc., Sunderland, MA, pp. 402–423.

Hess, G. R., 1994. Conservation Corridors and Contagious Disease: A Cautionary Note. Conservation Biology, 8(1), 256–262. http://www.jstor.org/stable/2386739

Horváth, Z., Ptacnik, R., Vad, C.F., Chase, J.M., 2019. Habitat loss over six decades accelerates regional and local biodiversity loss via changing landscape connectance. Ecol. Lett. 22, 1019–1027. 10.1111/ele.13260

Howeth, J.G., Leibold, M.A., 2013. Predation inhibits the positive effect of dispersal on intraspecific and interspecific synchrony in pond metacommunities. Ecology 94, 2220–2228. 10.1890/12-2066.1

Howeth, J.G., Leibold, M.A., 2010. Species dispersal rates alter diversity and ecosystem stability in pond metacommunities. Ecology 91, 2727–2741. 10.1890/09-1004.1

Hsieh, T.C., Ma, K.H., Chao, A., 2022. iNEXT: Interpolation and extrapolation for species diversity. R package version 3.0.0. http://chao.stat.nthu.edu.tw/wordpress/software_download/

Johnson, P.T.J., Hoverman, J.T., McKenzie, V.J., Blaustein, A.R., Richgels, K.L.D., 2013. Urbanization and wetland communities: applying metacommunity theory to understand the local and landscape effects. J. Appl. Ecol. 50, 34–42. 10.1111/1365-2664.12022

Joubert, L., Pryke, J.S., Samways, M.J., 2017. Moderate grazing sustains plant diversity in Afromontane grassland. Appl. Veg. Sci. 20, 340–351. 10.1111/avsc.12310

Kim, B.-J., Lee, B.-K., Lee, H., Jang, G.-S., 2016. Considering threats to population viability of the endangered Korean long-tailed goral (*Naemorhedus caudatus*) using VORTEX. Anim. Cells Syst. 20, 52–59. 10.1080/19768354.2015.1127856

Kindt, R., Coe, R., 2005. Tree diversity analysis: a manual and software for common statistical methods for ecological and biodiversity studies. World Agrofirestry Centre, Nairobi, Kenya.

Klinga, P., Mikoláš, M., Smolko, P., Tejkal, M., Höglund, J., Paule, L., 2019. Considering landscape connectivity and gene flow in the Anthropocene using complementary landscape genetics and habitat modelling approaches. Landsc. Ecol. 34, 521–536. 10.1007/s10980-019-00789-9

Lande, R., Engen, S., Sæther, B.E., 2003. Stochastic Population Dynamics in Ecology and Conservation. Oxford University Press, New York.

Leibold, M.A., Holyoak, M., Mouquet, N., Amarasekare, P., Chase, J.M., Hoopes, M.F., Holt, R.D., Shurin, J.B., Law, R., Tilman, D., Loreau, M., Gonzalez, A., 2004. The metacommunity concept: a framework for multiCscale community ecology. Ecol. Lett. 7, 601–613. 10.1111/j.1461-0248.2004.00608.x

Levin, S.A., Carpenter, S.R. (Eds.), 2012. The Princeton guide to ecology, 2. pr., 1. paperback pr. ed. Princeton University Press, Princeton, NJ.

Limberger, R., Pitt, A., Hahn, M.W., Wickham, S.A., 2019. Spatial insurance in multiCtrophic metacommunities. Ecol. Lett. 22, 1828–1837. 10.1111/ele.13365

Loeuille, N., Leibold, M.A., 2008. Evolution in Metacommunities: On the Relative Importance of Species Sorting and Monopolization in Structuring Communities. Am. Nat. 171, 788–799. 10.1086/587745

Lubchenco, J., 1978. Plant Species Diversity in a Marine Intertidal Community: Importance of Herbivore Food Preference and Algal Competitive Abilities. Am. Nat. 112, 23–39. 10.1086/283250

Luef, B., Aspetsberger, F., Hein, T., Huber, F., Peduzzi, P., 2007. Impact of hydrology on freeCliving and particleCassociated microorganisms in a river floodplain system (Danube, Austria). Freshw. Biol. 52, 1043–1057. 10.1111/j.1365-2427.2007.01752.x

Lüdecke, D., 2023. sjPlot: Data visualization for statistics in social science. R package version 2.8.15. https://CRAN.R-project.org/package=sjPlot

Magrach, A., Laurance, W.F., Larrinaga, A.R., Santamaria, L., 2014. MetaCAnalysis of the Effects of Forest Fragmentation on Interspecific Interactions. Conserv. Biol. 28, 1342–1348. 10.1111/cobi.12304

Magura, T., Ködöböcz, V., Tóthmérész, B., 2001. Effects of habitat fragmentation on carabids in forest patches. J. Biogeogr. 28, 129–138. 10.1046/j.1365-2699.2001.00534.x

Marini, L., Öckinger, E., Bergman, K., Jauker, B., Krauss, J., Kuussaari, M., Pöyry, J., Smith, H.G., SteffanCDewenter, I., Bommarco, R., 2014. Contrasting effects of habitat area and connectivity on evenness of pollinator communities. Ecography 37, 544–551. 10.1111/j.1600-0587.2013.00369.x

Matthies, D., Bräuer, I., Maibom, W., Tscharntke, T., 2004. Population size and the risk of local extinction: empirical evidence from rare plants. Oikos 105, 481–488. 10.1111/j.0030-1299.2004.12800.x

McCann, K.S., Rasmussen, J.B., Umbanhowar, J., 2005. The dynamics of spatially coupled food webs. Ecol. Lett. 8, 513–523. 10.1111/j.1461-0248.2005.00742.x

McGlinn, D.J., Xiao, X., May, F., Gotelli, N.J., Engel, T., Blowes, S.A., Knight, T.M., Purschke, O., Chase, J.M., McGill, B.J., 2019. Measurement of Biodiversity (MoB): A method to separate the scaleCdependent effects of species abundance distribution, density, and aggregation on diversity change. Methods Ecol. Evol. 10, 258–269. 10.1111/2041-210X.13102

Mony, C., Vandenkoornhuyse, P., Bohannan, B.J.M., Peay, K., Leibold, M.A., 2020. A Landscape of Opportunities for Microbial Ecology Research. Front. Microbiol. 11, 561427. 10.3389/fmicb.2020.561427

Murphy, M.V., 2022. semEff: Automatic Calculation of Effects for Piecewise Structural Equation Models. R package version 0.6.1. https://CRAN.R-project.org/package=semEff

Oertli, B., Parris, K.M., 2019. Review: Toward management of urban ponds for freshwater biodiversity. Ecosphere 10, e02810. 10.1002/ecs2.2810

Padial, A.A., Ceschin, F., Declerck, S.A.J., De Meester, L., Bonecker, C.C., Lansac-Tôha, F.A., Rodrigues, L., Rodrigues, L.C., Train, S., Velho, L.F.M., Bini, L.M., 2014. Dispersal Ability Determines the Role of Environmental, Spatial and Temporal Drivers of Metacommunity Structure. PLoS ONE 9, e111227. 10.1371/journal.pone.0111227

Pagano, M., 2008. Feeding of tropical cladocerans (Moina micrura, Diaphanosoma excisum) and rotifer (Brachionus calyciflorus) on natural phytoplankton: effect of phytoplankton size-structure. J. Plankton Res. 30, 401–414. 10.1093/plankt/fbn014

Peh, K.S.H., Lin YangChen, L.Y., Luke, S.H., Foster, W.A., Turner, E.C., 2014. Forest fragmentation and ecosystem function., in: Kettle, C.J., Koh, L.P. (Eds.), Global Forest Fragmentation. CABI, UK, pp. 96–114. 10.1079/9781780642031.0096

Pimm, S.L., 2008. Biodiversity: Climate Change or Habitat Loss — Which Will Kill More Species? Curr. Biol. 18, R117–R119. 10.1016/j.cub.2007.11.055

Pourriot R., 1977. Food and feeding habits of Rotifera. Archiv für Hydrobiologie-Beiheft Ergebnisse der Limnologie 8, 243–260.

Pulungan, M.A., Suzuki, S., Gavina, M.K.A., Tubay, J.M., Ito, H., Nii, M., Ichinose, G., Okabe, T., Ishida, A., Shiyomi, M., Togashi, T., Yoshimura, J., Morita, S., 2019. Grazing enhances species diversity in grassland communities. Sci. Rep. 9, 11201. 10.1038/s41598-019-47635-1

R Core Team, 2022. R: A language and environment for statistical computing. R Foundation for Statistical Computing, Vienna, Austria. https://www.R-project.org/.

Rösch, V., Tscharntke, T., Scherber, C., Batáry, P., 2013. Landscape composition, connectivity and fragment size drive effects of grassland fragmentation on insect communities. J. Appl. Ecol. 50, 387–394. 10.1111/1365-2664.12056

Rothhaupt, K.., 1990. Differences in particle size-dependent feeding efficiencies of closely related rotifer species. Limnol. Oceanogr. 35, 16–23. 10.4319/lo.1990.35.1.0016

Ryberg, W.A., Smith, K.G., Chase, J.M., 2012. Predators alter the scaling of diversity in prey metacommunities. Oikos 121, 1995–2000. 10.1111/j.1600-0706.2012.19620.x

Schloss, P.D., Westcott, S.L., Ryabin, T., Hall, J.R., Hartmann, M., Hollister, E.B., Lesniewski, R.A., Oakley, B.B., Parks, D.H., Robinson, C.J., Sahl, J.W., Stres, B., Thallinger, G.G., Van Horn, D.J., Weber, C.F., 2009. Introducing mothur: Open-Source, Platform-Independent, Community-Supported Software for Describing and Comparing Microbial Communities. Appl. Environ. Microbiol. 75, 7537–7541. 10.1128/AEM.01541-09

Shea, K., Roxburgh, S.H., Rauschert, E.S.J., 2004. Moving from pattern to process: coexistence mechanisms under intermediate disturbance regimes. Ecol. Lett. 7, 491–508. 10.1111/j.1461-0248.2004.00600.x

Simpson, G., 2023. gratia: Graceful ggplot-based graphics and other functions for GAMs fitted using mgcv. R package version 0.8.1. https://gavinsimpson.github.io/gratia/

Sinclair, J.S., Arnott, S.E., 2018. Local context and connectivity determine the response of zooplankton communities to salt contamination. Freshw. Biol. 63, 1273–1286. 10.1111/fwb.13132

Sterner, R.W., 1989. The role of grazers in phytoplankton succession. In Sommer, U. (ed.), Plankton Ecology. Springer-Verlag,Berlin: 107–170.

Szabó, B., Szabó, A., Vad, C.F., Boros, E., Lukić, D., Ptacnik, R., Márton, Z., Horváth, Z., 2022. Microbial stowaways: Waterbirds as dispersal vectors of aquatic proC and microeukaryotic communities. J. Biogeogr. jbi.14381. 10.1111/jbi.14381

Thompson, P., Forbes, C., Bernhardt, J., Davis, K., Stark, K., Yangel, E., Amadeo, F., Westwood, N., O’Connor, M., 2023. Biotic interactions structure zooplankton metacommunity dynamics following a summer heatwave (preprint). Preprints. 10.22541/au.167569427.75494960/v1

Thompson, P.L., Beisner, B.E., Gonzalez, A., 2015. Warming induces synchrony and destabilizes experimental pond zooplankton metacommunities. Oikos 124, 1171– 1180. 10.1111/oik.01945

Thompson, P.L., Guzman, L.M., De Meester, L., Horváth, Z., Ptacnik, R., Vanschoenwinkel, B., Viana, D.S., Chase, J.M., 2020. A processCbased metacommunity framework linking local and regional scale community ecology. Ecol. Lett. 23, 1314–1329. 10.1111/ele.13568

Thompson, P.L., Rayfield, B., Gonzalez, A., 2017. Loss of habitat and connectivity erodes species diversity, ecosystem functioning, and stability in metacommunity networks. Ecography 40, 98–108. 10.1111/ecog.02558

Thompson, P.L., Shurin, J.B., 2012. Regional zooplankton biodiversity provides limited buffering of pond ecosystems against climate change. J. Anim. Ecol. 81, 251–259. 10.1111/j.1365-2656.2011.01908.x

Trombulak, S.C., Frissell, C.A., 2000. Review of Ecological Effects of Roads on Terrestrial and Aquatic Communities. Conserv. Biol. 14, 18–30. 10.1046/j.1523-1739.2000.99084.x

Turunen, J., Louhi, P., Mykrä, H., Aroviita, J., Putkonen, E., Huusko, A., Muotka, T., 2018. Combined effects of local habitat, anthropogenic stress, and dispersal on stream ecosystems: a mesocosm experiment. Ecol. Appl. 28, 1606–1615. 10.1002/eap.1762

Urban, M.C., De Meester, L., 2009. Community monopolization: local adaptation enhances priority effects in an evolving metacommunity. Proc. R. Soc. B Biol. Sci. 276, 4129– 4138. 10.1098/rspb.2009.1382

Vad, C.F., HannyCEndrédi, A., Kratina, P., Abonyi, A., Mironova, E., Murray, D.S., Samchyshyna, L., Tsakalakis, I., Smeti, E., Spatharis, S., Tan, H., Preiler, C., Petrusek, A., Bengtsson, M.M., Ptacnik, R., 2023. Spatial insurance against a heatwave differs between trophic levels in experimental aquatic communities. Glob. Change Biol. 29, 3054–3071. 10.1111/gcb.16692

Vad, C.F., Péntek, A.L., Cozma, N.J., Földi, A., Tóth, A., Tóth, B., Böde, N.A., Móra, A., Ptacnik, R., Ács, É., Zsuga, K., Horváth, Z., 2017. Wartime scars or reservoirs of biodiversity? The value of bomb crater ponds in aquatic conservation. Biol. Conserv. 209, 253–262. 10.1016/j.biocon.2017.02.025

Valente, J.J., Gannon, D.G., Hightower, J., Kim, H., Leimberger, K.G., Macedo, R., Rousseau, J.S., Weldy, M.J., Zitomer, R.A., Fahrig, L., Fletcher, R.J., Wu, J., Betts, M.G., 2023. Toward conciliation in the habitat fragmentation and biodiversity debate. Landsc. Ecol. 38, 2717–2730. 10.1007/s10980-023-01708-9

Wardle, D.A., 2016. Do experiments exploring plant diversity–ecosystem functioning relationships inform how biodiversity loss impacts natural ecosystems? J. Veg. Sci. 27, 646–653. 10.1111/jvs.12399

Wardle, D.A., Zackrisson, O., 2005. Effects of species and functional group loss on island ecosystem properties. Nature 435, 806–810. 10.1038/nature03611

Williams, W.D., 2002. Environmental threats to salt lakes and the likely status of inland saline ecosystems in 2025. Environ. Conserv. 29, 154–167. 10.1017/S0376892902000103

Wood, S.N., 2017. Generalized Additive Models: An introduction with R (2nd edition). Chapman and Hall/CRC Press.

Woody, S.T., Ives, A.R., Nordheim, E.V., Andrews, J.H., 2007. DISPERSAL, DENSITY DEPENDENCE, AND POPULATION DYNAMICS OF A FUNGAL MICROBE ON LEAF SURFACES. Ecology 88, 1513–1524. 10.1890/05-2026

WWF, 2018. Living Planet Report - 2018: Aiming Higher. Grooten, M. and Almond, R.E.A.(Eds). WWF, Gland, Switzerland. LPR2018_Full_Report_Spreads.pdf (rackcdn.com)

Yuan, Z.Y., Jiao, F., Li, Y.H., Kallenbach, R.L., 2016. Anthropogenic disturbances are key to maintaining the biodiversity of grasslands. Sci. Rep. 6, 22132. 10.1038/srep22132

Zubkov, M.V., 2014. Faster growth of the major prokaryotic versus eukaryotic CO2 fixers in the oligotrophic ocean. Nat. Commun. 5, 3776. 10.1038/ncomms4776

