## Supplementary material for "Connectivity loss in experimental pond networks leads to biodiversity loss in microbial communities"

This file includes the following text, tables and figures:

Text S1-S3

Table S1, Tables S3-S4, Table S6, Table S11-S19

Figures S1-S9

The following other supplementary materials for this manuscript are available as separate Excel spreadsheets:

Tables S2, S5, Tables S7-S10, Table S20

Text S1. Description of inoculum creation.

To create the inoculum, 10 plastic barrels were set up, the same type as those used for the mesocosms, and filled with 200 liters of tap water. The barrels were covered with mosquito nets and allowed to stand for four days, from June 26^th^ to June 30^th^, to allow for de-chlorination through evaporation. Next, equal amounts of sediment were added to the mesocosms, collected from 10 intermittent eutrophic pools and ponds (Table S1). Each of the 10 mesocosms received sediment from one pond. After the addition of the sediment, the water was mixed, and a six-day hatching period was initiated, ending on July 6^th^. During this period, water in the mesocosms was mixed thoroughly each day.

At the end of the hatching period, water of the 10 mesocosms were pooled to create the inoculum. Before distributing this pooled community to the experimental mesocosms, 21 L of water was removed from each mesocosm to make space for the inoculum water, then 21 L of inoculum water was distributed to each individual experimental mesocosm resulting in a total volume of 200 L of water at the start of the experiment.

Text S2. Detailed description for sampling and identification of phytoplankton and zooplankton communities, for measurement of chlorophyll-a fluorescence, and determination of physical and chemical variables.

*Phytoplankton communities*

The community composition of nano- and microplankton was determined with an inverted microscope (Zeiss Axiovert) (Utermöhl, 1958) at 630x magnification using the relevant taxonomic guide (John et al., 2002). As the total number of enumerated individuals largely differed across the samples, 15 samples with more than 400 individuals were rarefied to 400 counts prior to analyses on species richness. In the other 15 samples, where it was not possible to enumerate up to 400 individuals due to the low phytoplankton density, no rarefaction was applied.

*Zooplankton communities*

Individuals of zooplankton were identified to genus level within the three main groups of zooplankton (rotifers, cladocerans, ostracods) according to the relevant identification guides (Bancsi, 1986; Gulyás and Forró, 1999). Crustacean groups (Cladocera and Ostracoda) were identified with Olympus SZ51 at 0.8x-4x magnification, while Rotifera with Olympus CX33 upright microscope at 4x-100x magnification. To enumerate Crustacean and Rotifera, subsampling was applied if the density was too high to identify all individuals in the whole sample (Herzig, 1984). Per sample, 200 individuals were enumerated.

Dry weight was calculated for each taxon (hereinafter referred to as zooplankton biomass - ZB, µg L^-1^) based on length-weight relationships (Bottrell et al. 1976) using body length data from relevant literature in the case of both Crustacean and Rotifera (Bottrell et al. 1976; SOP LG403, 2016).

When estimating zooplankton taxon richness, bdelloid rotifers (Bdelloidea) were excluded since it was not possible to identify them to genus level. For the analyses on zooplankton biomass, all taxa were included.

*Chlorophyll-a fluorescence*

15 ml of composite water sample was taken from three different surface spots of each mesocosm in the morning of the sampling day. After a 30-minute dark adaptation period to avoid potential bias resulting from short-term physiological changes (Perri et al., 2021), chlorophyll-a fluorescence (maximum fluorescence yield, Chl-a, a.u.) measurement was carried out on 455 nm wavelength with a handheld fluorometer (AquaPen AP 110-C, Photon System Instruments) using the OJIP protocol (Stirbet and Govindjee, 2011).

*Physical and chemical variables*

Temperature (°C), pH and electric conductivity (EC, μS cm^-1^) were measured *in situ* using HANNA HI-9142 and HI-98129 field devices (HANNA Instruments, USA).

To determine nutrients, a total of 1.5 L of water was collected with a plexiglass tube sampler from three different locations (3 x 0.5 L) of each mesocosm after thorough mixing and filtered through a 100-μm mesh plankton net. For further processing, 200 mL of the composite sieved water was immediately delivered to the laboratory in a glass bottle in a cool box. Total nitrogen (TN, mg L^-1^) concentration was determined with a Multi N/C 3100 TC/TN analyzer (Analytik Jena, Germany) and total phosphorus (TP, μg L^-1^) concentration with standard spectrophotometric method from unfiltered samples followed by persulphate acidic digestion (Eaton et al., 2005).

Text S3. Detailed description of DNA isolation, amplification, sequencing and amplicon data analysis.

*DNA isolation, amplification and sequencing*

DNA extraction from the membrane filters was performed using the DNeasy PowerSoil Kit (Qiagen, Germany), according to the manufacturer's instructions, with the following exceptions: cell disruption was performed using an MM301 mixer mill (Retsch, Germany) with shaking for 2 min at 30 Hz, and 50 μL of C6 solution was used instead of the 100 μL suggested. The extracted DNA was stored at -20 °C.

The isolated DNA was amplified using polymerase chain reaction (PCR) to amplify fragments of taxonomic marker genes encoding 16S and 18S ribosomal RNA. The primers used for the reaction were 574*F (CGGTAAYTCCAGCTCYAV) and 1132R (CCGTCAATTHCTTYAART) (Hugerth et al., 2014) in the case of eukaryotes, and primers 341F (CCTACGGGGGNGGCWGCAG) (Herlemann et al., 2011) and 805NR (GACTACHVGGGTATCTAATCC) (Apprill et al., 2015) were used to identify bacteria.

The PCR was performed according to the following protocol: the final volume of the total reaction mixture was 20 μL, containing 1 × Phusion HF Buffer (Thermo Fischer Scientific, USA), 0.2 mM dinucleotide triphosphate mixture (dNTPs), 10 μL of RNase and DNase free DEPC treated sterile water, 0.325 µM of each primer, 0.4 U Phusion High-Fidelity DNA Polymerase (Thermo Fisher Scientific, USA), 0.4 µg µL^-1^ Bovine Serum Albumin (Fermentas) and 1 μL of DNA sample. For proper amplification, we deviated from the above-described mixture in the case of eukaryotes and some bacteria samples by adding 2 or 3 μL of sample DNA to the reaction mixture and reducing the water to reach the same, 20 μL total volume. The reaction temperature profile was adjusted to be specific for the 18S rRNA gene as follows: initial denaturation 5 min at 98 °C, followed by 25 cycles of 95 °C for 30 s, 51 °C for 30 s, 72 °C for 1 min, and a final extension of 72 °C for 10 min. For amplification specific to the 16S rRNA gene, annellation was carried out at 55 °C for 30 s.

The PCR products were checked by agarose gel electrophoresis. We were not able to successfully amplify sequences from the sample of one of the mesocosms belonging to a connected metacommunity (M3) at week 12.

The PCR products were sent to the Genomics Core Facility RTSF, Michigan State University, USA for sequencing using MiSeq v2 reagent kit in 2×250 bp format on Illumina MiSeq platform (Illumina, USA).

*Amplicon data processing*

The DNA sequences were then processed with mothur v.1.47.0 (Schloss et al., 2009) following the MiSeq SOP (<http://www.mothur.org/wiki/MiSeq_SOP>; Kozich et al., 2013; downloaded on 22^nd^ March 2022) with some modifications. In the "make.contigs'' command, we set the deltaq value to 10. We used the “trim.seqs” command, setting the checkorient parameter to true, to remove the primers and their possible reverse complements from the sequences. In the "pre.cluster" command we allowed 4 differences between sequences, in the "split.abund" command we removed singleton sequences after chimera filtering. Thereafter, we followed the SOP to identify amplicon sequence variants (ASVs). For the classification of bacteria after finishing the procedure with mothur we used TaxAss (Rohwer et al., 2018) with ARB-SILVA SSU v138.1 (Quast et al., 2012) and FreshTrain (15^th^ June 2020 release; Newton et al., 2011) reference databases for a more precise ASV classification. In the case of eukaryotes, we differed from the MiSeq SOP by using the PR2 database (Guillou et al., 2012) instead of the ARB-SILVA database for a better classification. ASVs assigned to “Archaea”, “Chloroplast”, “Mitochondria” and “unknown” groups were excluded from the analysis. In the case of eukaryotes, ASVs assigned to “Metazoa”, “Basidiomycota”, “Ascomycota”, “Zygnemophyceae_X_unclassified”, “Zygnemophyceae_XX_unclassified”, “Ulmus”, “Spirogyra”, “Cylindrocystis”, “Streptophyta_unclassified”, “Zannichellia”, “Taxus”, “Embryophyceae_XX_unclassified”, “Juncus”, “Secale”, “Allium”, “Helianthus”, “Brachypodium” were removed from the analysis. The latter removal was deemed to be necessary because sequences assigned to these taxa were most probably present in the samples due to contamination or they represented taxa that might have caused an algal bloom in the mesocosms. After the taxonomic identification, we decided to not include the eukaryotes from one of the mesocosms within a control metacommunity (M1) at the first sampling, as the number of identified sequences were less than 2000, and this could have affected our overall analysis.

References for Text S1, S2, S3

Apprill, A., McNally, S., Parsons, R., & Weber, L. (2015). Minor revision to V4 region SSU rRNA 806R gene primer greatly increases detection of SAR11 bacterioplankton. *Aquatic Microbial Ecology*, *75*(2), 129–137. https://doi.org/10.3354/ame01753

Bancsi, I. (1986). A kerekesférgek (Rotatoria) kishatározója I. Vizügyi Hidrobiológia 15. Vízgazdálkodási Intézet, Budapest, 1–172.

Bottrell, H. H., Duncan, A., Gliwicz, Z. M., Grygierek, E., Herzig, A., Hillbricht-Ilkowska, A., Kurasawa, H., Larsson, P., & Weglenska, T. (1976). A review of some problems in zooplankton production studies. *Norwegian Journal of Zoology*, *24*, 419-456.

Eaton, A. D., Clesceri, L. S., Rice, E. W., Greenberg, A. E., & Franson, M. A. H. (eds.) (2005). Standard Methods for the Examination of Water & Wastewater, Centennial Edition (21^st^ ed). American Public Health Association (APHA) Press, Washington, DC.

Guillou, L., Bachar, D., Audic, S., Bass, D., Berney, C., Bittner, L., Boutte, C., Burgaud, G., de Vargas, C., Decelle, J., del Campo, J., Dolan, J. R., Dunthorn, M., Edvardsen, B., Holzmann, M., Kooistra, W. H. C. F., Lara, E., Le Bescot, N., Logares, R., Mahé, F., Massana, R., Montresor, M., Morard, R., Not, F., Pawlowski, J., Probert, I., Sauvadet, A.-L., Siano, R., Stoeck, T., Vaulot, D., Zimmermann, P., & Christen, R. (2012). The Protist Ribosomal Reference database (PR2): a catalog of unicellular eukaryote Small Sub-Unit rRNA sequences with curated taxonomy. *Nucleic Acids Research*, *41*(D1), D597–D604. https://doi.org/10.1093/nar/gks1160

Gulyás, P., & Forró, L. (1999). Az ágascsápú rákok (Cladocera) kishatározója. *Vízi természet- és környezetvédelem*, *9*, 1-237.

Herlemann, D. P., Labrenz, M., Jürgens, K., Bertilsson, S., Waniek, J. J., & Andersson, A. F. (2011). Transitions in bacterial communities along the 2000 km salinity gradient of the Baltic Sea. *The ISME Journal*, *5*(10), 1571–1579. https://doi.org/10.1038/ismej.2011.41

Herzig, A. (1984). Fundamental requirements for zooplankton production studies. Institute for Limnology of the Austrian Academy of Sciences, Mondsee.

John, D. M., Whitton, B. A., & Brook, A. J., (eds.) (2002). The freshwater algal flora of the British Isles: An identification guide to freshwater and terrestrial algae. Cambridge University Press.

Kozich, J. J., Westcott, S. L., Baxter, N. T., Highlander, S. K., & Schloss, P. D. (2013). Development of a Dual-Index Sequencing Strategy and Curation Pipeline for Analyzing Amplicon Sequence Data on the MiSeq Illumina Sequencing Platform. *Applied and Environmental Microbiology*, *79*(17), 5112–5120. https://doi.org/10.1128/AEM.01043-13

Newton, R. J., Jones, S. E., Eiler, A., McMahon, K. D., & Bertilsson, S. (2011). A Guide to the Natural History of Freshwater Lake Bacteria. *Microbiology and Molecular Biology Reviews*, *75*(1), 14–49. https://doi.org/10.1128/MMBR.00028-10

Perri, K. A., Manning, S. R., Watson, S. B., Fowler, N. L., & Boyer, G. L. (2021). Dark adaptation and ability of pulse-amplitude modulated (PAM) fluorometry to identify nutrient limitation in the bloom-forming cyanobacterium, *Microcystis aeruginosa* (Kützing). *Journal of Photochemistry and Photobiology B: Biology*, *219*, 112186. https://doi.org/10.1016/j.jphotobiol.2021.112186

Quast, C., Pruesse, E., Yilmaz, P., Gerken, J., Schweer, T., Yarza, P., Peplies, J., & Glöckner, F. O. (2012). The SILVA ribosomal RNA gene database project: improved data processing and web-based tools. *Nucleic Acids Research*, *41*(D1), D590–D596. https://doi.org/10.1093/nar/gks1219

Rohwer, R. R., Hamilton, J. J., Newton, R. J., & McMahon, K. D. (2018). TaxAss: Leveraging a Custom Freshwater Database Achieves Fine-Scale Taxonomic Resolution. *mSphere*, *3*(5), e00327-18. https://doi.org/10.1128/mSphere.00327-18

Schloss, P. D., Westcott, S. L., Ryabin, T., Hall, J. R., Hartmann, M., Hollister, E. B., Lesniewski, R. A., Oakley, B. B., Parks, D. H., Robinson, C. J., Sahl, J. W., Stres, B., Thallinger, G. G., Van Horn, D. J., & Weber, C. F. (2009). Introducing mothur: Open-Source, Platform-Independent, Community-Supported Software for Describing and Comparing Microbial Communities. *Applied and Environmental Microbiology*, *75*(23), 7537–7541. https://doi.org/10.1128/AEM.01541-09

SOP LG403 (2016). Standard Operating Procedure for Zooplankton Analysis. Revision 07, July 2016. Great Lakes National Program Office, U.S. Environmental Protection Agency, Chicaco, IL.

Stirbet, A., & Govindjee (2011). On the relation between the Kautsky effect (chlorophyll a fluorescence induction) and Photosystem II: Basics and applications of the OJIP fluorescence transient. *Journal of Photochemistry and Photobiology B: Biology*, *104*(1-2), 236–257. https://doi.org/10.1016/j.jphotobiol.2010.12.010

Utermöhl, H. (1958). Zur Vervollkommnung der quantitativen Phytoplankton-Methodik. *Mitteilung Internationale Vereinigung Fuer Theoretische unde Amgewandte Limnologie*, *9*, 1–38.

Table S1. GPS coordinates of the soda pans, bomb craters, and a temporary pool where sediment samples (top 3 cm layer from the dry sediment) used for inoculation were collected.

| **Name** | **Latitude** | **Longitude** |
| --- | --- | --- |
| Albersee | 47.77513639 | 16.77011917 |
| Kirchsee | 47.75867861 | 16.78548861 |
| Krautingsee | 47.75602861 | 16.78047972 |
| Martenhofenlacken | 47.75050167 | 16.85669500 |
| Obere Höllacke | 47.82711944 | 16.80793222 |
| Oberer Stinkersee | 47.81376222 | 16.79251722 |
| Westliche Wörthenlacke | 47.77092167 | 16.87077667 |
| Bomb crater #1 | 47.12111100 | 19.13344400 |
| Bomb crater #2 | 47.12208974 | 19.13614305 |
| Wheel track pool | 47.12130600 | 19.13169400 |

Table S3. Summary table of nested ANOVA performed on filamentous algae biomass (FB, g L^-1^) in week 8, 12 and 16. Applied data transformation is presented along with the abbreviation of each variable.

|  |  |  | Df | Sum Sq | Mean Sq | F-value | P-value |
| --- | --- | --- | --- | --- | --- | --- | --- |
| **WEEK 8** | FB^^0.25^ | Treatment | 1 | 0.0004 | 0.0004 | 0.012 | 0.9150 |
|  |  | Treatment:Meta | 4 | 0.2042 | 0.0510 | 1.483 | 0.2430 |
|  |  | Residuals | 21 | 0.7230 | 0.0344 |  |  |
| **WEEK 12** | FB^^0.25^ | Treatment | 1 | 0.0151 | 0.0151 | 0.476 | 0.4970 |
|  |  | Treatment:Meta | 4 | 0.2669 | 0.0667 | 2.104 | 0.1120 |
|  |  | Residuals | 24 | 0.7610 | 0.0317 |  |  |
| **WEEK 16** | FB^^0.333^ | Treatment | 1 | 0.0025 | 0.0025 | 0.065 | 0.8020 |
|  |  | Treatment:Meta | 4 | 0.2434 | 0.0608 | 1.568 | 0.2150 |
|  |  | Residuals | 24 | 0.9310 | 0.0388 |  |  |

Table S4. Summary table of nested PERMANOVA performed on z-score standardized environmental variables (Euclidean distance; permutations=2000).

|  |  | Df | Sum Sq | F-value | P-value |  |
| --- | --- | --- | --- | --- | --- | --- |
| **WEEK 4** | Treatment | 1 | 1.934 | 0.524 | 0.7981 |  |
|  | Treatment:Meta | 4 | 14.763 | 0.690 | 0.8156 |  |
|  | Residual | 24 | 128.304 |  |  |  |
| **WEEK 8** | Treatment | 1 | 5.868 | 1.885 | 0.2054 |  |
|  | Treatment:Meta | 4 | 12.450 | 0.590 | 0.9145 |  |
|  | Residual | 24 | 126.683 |  |  |  |
| **WEEK 12** | Treatment | 1 | 8.800 | 1.268 | 0.4108 |  |
|  | Treatment:Meta | 4 | 27.766 | 1.536 | 0.0910 | . |
|  | Residual | 24 | 108.434 |  |  |  |
| **WEEK 16** | Treatment | 1 | 7.113 | 1.603 | 0.2994 |  |
|  | Treatment:Meta | 4 | 17.746 | 0.886 | 0.5962 |  |
|  | Residual | 24 | 120.140 |  |  |  |

Table S6. Summary table of nested ANOVA performed on the taxon richness (^α^S) and biomass (ZB, µg L^-1^) at α-scale in zooplankton. Applied data transformation is presented along with the abbreviation of each variable.

|  | **Richness** | | | | | | | |
| --- | --- | --- | --- | --- | --- | --- | --- | --- |
|  |  |  | Df | Sum Sq | Mean Sq | F-value | P-value |  |
| **WEEK 4** | ^α^S | Treatment | 1 | 0.300 | 0.3000 | 0.273 | 0.6063 |  |
|  |  | Treatment:Meta | 4 | 10.670 | 2.6670 | 2.424 | 0.0759 | . |
|  |  | Residuals | 24 | 26.400 | 1.1000 |  |  |  |
| **WEEK 8** | ^α^S | Treatment | 1 | 0.530 | 0.5333 | 0.320 | 0.5770 |  |
|  |  | Treatment:Meta | 4 | 1.330 | 0.3333 | 0.200 | 0.9360 |  |
|  |  | Residuals | 24 | 40.000 | 1.6667 |  |  |  |
| **WEEK 12** | ^α^S | Treatment | 1 | 0.130 | 0.133 | 0.072 | 0.7906 |  |
|  |  | Treatment:Meta | 4 | 42.930 | 10.733 | 5.802 | **0.0021** | ***** |
|  |  | Residuals | 24 | 44.400 | 1.8500 |  |  |  |
| **WEEK 16** | ^α^S^^0.25^ | Treatment | 1 | 0.0005 | 0.0005 | 0.046 | 0.8318 |  |
|  |  | Treatment:Meta | 4 | 0.1538 | 0.0384 | 3.885 | **0.0143** | ***** |
|  |  | Residuals | 24 | 0.2375 | 0.0099 |  |  |  |
|  | **Biomass** | | | | | | | |
|  |  |  | Df | Sum Sq | Mean Sq | F-value | P-value |  |
| **WEEK 4** | log(ZB) | Treatment | 1 | 0.168 | 0.1679 | 0.383 | 0.5417 |  |
|  |  | Treatment:Meta | 4 | 0.607 | 0.1517 | 0.346 | 0.8440 |  |
|  |  | Residuals | 24 | 10.512 | 0.4380 |  |  |  |
| **WEEK 8** | log(ZB) | Treatment | 1 | 0.651 | 0.6513 | 1.247 | 0.2752 |  |
|  |  | Treatment:Meta | 4 | 6.622 | 1.6555 | 3.170 | **0.0317** | ***** |
|  |  | Residuals | 24 | 12.535 | 0.5223 |  |  |  |
| **WEEK 12** | log(ZB) | Treatment | 1 | 4.024 | 4.0240 | 4.086 | 0.0545 | . |
|  |  | Treatment:Meta | 4 | 7.341 | 1.8350 | 1.864 | 0.1497 |  |
|  |  | Residuals | 24 | 23.634 | 0.9850 |  |  |  |
| **WEEK 16** | log(ZB) | Treatment | 1 | 7.160 | 7.1600 | 8.073 | **0.0090** | ****** |
|  |  | Treatment:Meta | 4 | 8.017 | 2.0040 | 2.260 | 0.0925 | . |
|  |  | Residuals | 24 | 21.287 | 0.8870 |  |  |  |

Table S11. Summary table of structural equation models (SEMs) showing the relationships (direct and indirect effects) between treatment, zooplankton biomass (ZB), and diversity (^α^S or J) in prokaryotes at the end of the experiment (week 16). Applied data transformation is presented along with the abbreviation of each variable. In the models, treatment and log(ZB) were included as fixed, and metacommunity ID as a nested random effect factor. Effects for which the confidence intervals do not contain zero are highlighted with a star (i.e., significant at the CI level).

| **Prokaryotes** | | | | | | | | |
| --- | --- | --- | --- | --- | --- | --- | --- | --- |
|  |  |  |  | Std. Effect | Bias | Std. Error | CI (95%) |  |
| SEM1 | log(ZB) | Direct | Treatment[fragm.] | -0.443 | 0.032 | 0.150 | -0.748 – -0.123 | ***** |
|  |  | Total | Treatment[fragm.] | -0.443 | 0.032 | 0.150 | -0.748 – -0.123 | ***** |
|  | ^α^S | Direct | Treatment[fragm.] | -0.187 | -0.014 | 0.183 | -0.483 – 0.214 |  |
|  |  |  | log(ZB) | 0.009 | -0.056 | 0.260 | -0.561 – 0.514 |  |
|  |  | Indirect | Treatment[fragm.] | -0.004 | 0.009 | 0.116 | -0.338 – 0.199 |  |
|  |  | Total | Treatment[fragm.] | -0.191 | -0.004 | 0.132 | -0.427 – 0.064 |  |
|  |  |  | log(ZB) | 0.009 | -0.056 | 0.260 | -0.561 – 0.514 |  |
|  |  | Mediators | log(ZB) | -0.004 | 0.009 | 0.116 | -0.338 – 0.199 |  |
| SEM2 | log(ZB) | Direct | Treatment[fragm.] | -0.443 | 0.032 | 0.150 | -0.748 – -0.123 | ***** |
|  |  | Total | Treatment[fragm.] | -0.443 | 0.032 | 0.150 | -0.748 – -0.123 | ***** |
|  | log(J) | Direct | Treatment[fragm.] | -0.084 | -0.034 | 0.222 | -0.519 – 0.287 |  |
|  |  |  | log(ZB) | -0.194 | -0.038 | 0.150 | -0.493 – 0.102 |  |
|  |  | Indirect | Treatment[fragm.] | 0.086 | 0.007 | 0.077 | -0.004 – 0.318 |  |
|  |  | Total | Treatment[fragm.] | 0.002 | -0.027 | 0.188 | -0.429 – 0.338 |  |
|  |  |  | log(ZB) | -0.194 | -0.038 | 0.150 | -0.493 – 0.102 |  |
|  |  | Mediators | log(ZB) | 0.086 | 0.007 | 0.077 | -0.004 – 0.318 |  |

Table S11. Summary table of structural equation models (SEMs) showing the relationships (direct and indirect effects) between treatment, zooplankton biomass (ZB), and diversity (^α^S or J) in microeukaryotes at the end of the experiment (week 16). Applied data transformation is presented along with the abbreviation of each variable. In the models, treatment and log(ZB) were included as fixed, and metacommunity ID as a nested random effect factor. Effects for which the confidence intervals do not contain zero are highlighted with a star (i.e., significant at the CI level). (continued)

| **Microeukaryotes** | | | | | | | | |
| --- | --- | --- | --- | --- | --- | --- | --- | --- |
|  |  |  |  | Std. Effect | Bias | Std. Error | CI (95%) |  |
| SEM1 | log(ZB) | Direct | Treatment[fragm.] | -0.443 | 0.032 | 0.150 | -0.748 – -0.123 | ***** |
|  |  | Total | Treatment[fragm.] | -0.443 | 0.032 | 0.150 | -0.748 – -0.123 | ***** |
|  | log(^α^S) | Direct | Treatment[fragm.] | -0.464 | -0.021 | 0.118 | -0.657 – -0.144 | ***** |
|  |  |  | log(ZB) | 0.248 | -0.085 | 0.175 | 0.019 – 0.665 | ***** |
|  |  | Indirect | Treatment[fragm.] | -0.110 | 0.038 | 0.063 | -0.269 – -0.011 | ***** |
|  |  | Total | Treatment[fragm.] | -0.574 | 0.017 | 0.096 | -0.722 – -0.365 | ***** |
|  |  |  | log(ZB) | 0.248 | -0.085 | 0.175 | 0.019 – 0.665 | ***** |
|  |  | Mediators | log(ZB) | -0.110 | 0.038 | 0.063 | -0.269 – -0.011 | ***** |
| SEM2 | log(ZB) | Direct | Treatment[fragm.] | -0.443 | 0.032 | 0.150 | -0.748 – -0.123 | ***** |
|  |  | Total | Treatment[fragm.] | -0.443 | 0.032 | 0.150 | -0.748 – -0.123 | ***** |
|  | log(J) | Direct | Treatment[fragm.] | -0.228 | -0.018 | 0.086 | -0.415 – -0.078 | ***** |
|  |  |  | log(ZB) | 0.476 | -0.063 | 0.149 | 0.201 – 0.674 | ***** |
|  |  | Indirect | Treatment[fragm.] | -0.211 | 0.044 | 0.073 | -0.364 – -0.094 | ***** |
|  |  | Total | Treatment[fragm.] | -0.439 | 0.026 | 0.096 | -0.604 – -0.232 | ***** |
|  |  |  | log(ZB) | 0.476 | -0.063 | 0.149 | 0.201 – 0.674 | ***** |
|  |  | Mediators | log(ZB) | -0.211 | -0.044 | 0.073 | -0.364 – -0.094 | ***** |

| **Prokaryotes** | | | | | | | | |
| --- | --- | --- | --- | --- | --- | --- | --- | --- |
|  |  |  |  | Std. Effect | Bias | Std. Error | CI (95%) |  |
| SEM1 | ^α^S | Direct | Treatment[fragm.] | -0.213 | -0.006 | 0.154 | -0.517 – 0.052 |  |
|  |  | Total | Treatment[fragm.] | -0.213 | -0.006 | 0.154 | -0.517 – 0.052 |  |
|  | log(ZB) | Direct | Treatment[fragm.] | -0.444 | 0.022 | 0.140 | -0.690 – -0.095 | ***** |
|  |  |  | ^α^S | -0.025 | -0.046 | 0.230 | -0.532 – 0.333 |  |
|  |  | Indirect | Treatment[fragm.] | 0.005 | 0.012 | 0.060 | -0.108 – 0.139 |  |
|  |  | Total | Treatment[fragm.] | -0.438 | 0.034 | 0.141 | -0.643 – -0.114 | ***** |
|  |  |  | ^α^S | -0.025 | -0.046 | 0.230 | -0.532 – 0.333 |  |
|  |  | Mediators | ^α^S | 0.005 | 0.012 | 0.060 | -0.108 – 0.139 |  |
| SEM2 | log(J) | Direct | Treatment[fragm.] | 0.002 | -0.032 | 0.208 | -0.490 – 0.370 |  |
|  |  | Total | Treatment[fragm.] | 0.002 | -0.032 | 0.208 | -0.490 – 0.370 |  |
|  | log(ZB) | Direct | Treatment[fragm.] | -0.443 | 0.027 | 0.167 | -0.747 – -0.084 | ***** |
|  |  |  | log(J) | -0.173 | -0.021 | 0.118 | -0.403 – 0.053 |  |
|  |  | Indirect | Treatment[fragm.] | 0.000 | 0.008 | 0.045 | -0.086 – 0.096 |  |
|  |  | Total | Treatment[fragm.] | -0.443 | 0.035 | 0.149 | -0.736 – -0.117 | ***** |
|  |  |  | log(J) | -0.173 | -0.021 | 0.118 | -0.403 – 0.053 |  |
|  |  | Mediators | log(J) | 0.000 | 0.008 | 0.045 | -0.086 – 0.096 |  |

Table S12. Summary table of structural equation models (SEMs) showing the relationships (direct and indirect effects) between treatment, zooplankton biomass (ZB), and diversity (^α^S or J) in prokaryotes at the end of the experiment (week 16). Applied data transformation is presented along with the abbreviation of each variable. In the models, treatment ^α^S and log(J) were included as fixed, and metacommunity ID as a nested random effect factor. Effects for which the confidence intervals do not contain zero are highlighted with a star (i.e., significant at the CI level).

Table S12. Summary table of structural equation models (SEMs) showing the relationships (direct and indirect effects) between treatment, zooplankton biomass (ZB), and diversity (^α^S or J) in microeukaryotes at the end of the experiment (week 16). Applied data transformation is presented along with the abbreviation of each variable. In the models, treatment, log(^α^S) and log(J) were included as fixed, and metacommunity ID as a nested random effect factor. Effects for which the confidence intervals do not contain zero are highlighted with a star (i.e., significant at the CI level). (continued)

| **Microeukaryotes** | | | | | | | | |
| --- | --- | --- | --- | --- | --- | --- | --- | --- |
|  |  |  |  | Std. Effect | Bias | Std. Error | CI (95%) |  |
| SEM1 | log(^α^S) | Direct | Treatment[fragm.] | -0.612 | 0.018 | 0.119 | -0.859 – -0.404 | ***** |
|  |  | Total | Treatment[fragm.] | -0.612 | 0.018 | 0.119 | -0.859 – -0.404 | ***** |
|  | log(ZB) | Direct | Treatment[fragm.] | -0.128 | -0.083 | 0.120 | -0.279 – 0.045 |  |
|  |  |  | log(^α^S) | 0.363 | -0.156 | 0.202 | 0.151 – 0.661 | ***** |
|  |  | Indirect | Treatment[fragm.] | -0.222 | 0.100 | 0.109 | -0.376 – -0.122 | ***** |
|  |  | Total | Treatment[fragm.] | -0.350 | 0.016 | 0.112 | -0.548 – -0.093 | ***** |
|  |  |  | log(^α^S) | 0.363 | -0.156 | 0.202 | 0.151 – 0.661 | ***** |
|  |  | Mediators | log(^α^S) | -0.222 | 0.100 | 0.109 | -0.376 – -0.122 |  |
| SEM2 | log(J) | Direct | Treatment[fragm.] | -0.489 | 0.027 | 0.132 | -0.752 – -0.239 | ***** |
|  |  | Total | Treatment[fragm.] | -0.489 | 0.027 | 0.132 | -0.752 – -0.239 | ***** |
|  | log(ZB) | Direct | Treatment[fragm.] | -0.147 | -0.030 | 0.107 | -0.383 – 0.029 |  |
|  |  |  | log(J) | 0.489 | -0.083 | 0.148 | 0.233 – 0.675 | ***** |
|  |  | Indirect | Treatment[fragm.] | -0.239 | 0.054 | 0.078 | -0.468 – -0.124 | ***** |
|  |  | Total | Treatment[fragm.] | -0.386 | 0.023 | 0.117 | -0.553 – -0.091 | ***** |
|  |  |  | log(J) | 0.489 | -0.083 | 0.148 | 0.232 – 0.675 | ***** |
|  |  | Mediators | log(J) | -0.239 | 0.054 | 0.078 | -0.468 – -0.124 | ***** |

Table S13. Summary table of nested ANOVA performed on the number of observed ASVs (^α^S), effective number of ASVs of PIE (^α^S_PIE_) and evenness (J) at α-scale, and Whittaker’s β-diversity (β_S_) in prokaryotes and microeukaryotes. Applied data transformation is presented along with the abbreviation of each variable. Significant P-values are marked in bold.

|  | **Prokaryotes** | | | | | | | | **Microeukaryotes** | | | | | | | |
| --- | --- | --- | --- | --- | --- | --- | --- | --- | --- | --- | --- | --- | --- | --- | --- | --- |
|  |  |  | Df | Sum Sq | Mean Sq | F-value | P-value |  |  |  | Df | Sum Sq | Mean Sq | F-value | P-value |  |
| **WEEK 4** | log(^α^S) | Treatment | 1 | 0.0093 | 0.0093 | 0.453 | 0.5070 |  | log(^α^S) | Treatment | 1 | 0.0124 | 0.0124 | 0.163 | 0.6900 |  |
|  |  | Treatment:Meta | 4 | 0.0782 | 0.0195 | 0.955 | 0.4500 |  |  | Treatment:Meta | 4 | 0.0848 | 0.0212 | 0.278 | 0.8890 |  |
|  |  | Residuals | 24 | 0.4912 | 0.0205 |  |  |  |  | Residuals | 23 | 1.7499 | 0.0761 |  |  |  |
| **WEEK 8** | ^α^S | Treatment | 1 | 208 | 208 | 0.110 | 0.7430 |  | log(^α^S) | Treatment | 1 | 0.0000 | 0.0000 | 0.002 | 0.9680 |  |
|  |  | Treatment:Meta | 4 | 7059 | 1765 | 0.930 | 0.4630 |  |  | Treatment:Meta | 4 | 0.1500 | 0.0375 | 1.605 | 0.2050 |  |
|  |  | Residuals | 24 | 45541 | 1898 |  |  |  |  | Residuals | 24 | 0.5607 | 0.0234 |  |  |  |
| **WEEK 12** | log(^α^S) | Treatment | 1 | 0.1365 | 0.1365 | 5.183 | **0.0324** | ***** | ^α^S | Treatment | 1 | 476 | 475.7 | 1.796 | 0.1930 |  |
|  |  | Treatment:Meta | 4 | 0.4265 | 0.1066 | 4.049 | **0.0125** | ***** |  | Treatment:Meta | 4 | 773 | 193.2 | 0.730 | 0.5810 |  |
|  |  | Residuals | 23 | 0.6056 | 0.0263 |  |  |  |  | Residuals | 23 | 6090 | 264.8 |  |  |  |
| **WEEK 16** | ^α^S | Treatment | 1 | 3162 | 3162 | 1.269 | 0.2710 |  | log(^α^S) | Treatment | 1 | 0.9882 | 0.9882 | 22.614 | **0.0001** | ******* |
|  |  | Treatment:Meta | 4 | 6924 | 1731 | 0.694 | 0.6030 |  |  | Treatment:Meta | 4 | 0.6001 | 0.1500 | 3.433 | **0.0235** | ***** |
|  |  | Residuals | 24 | 59818 | 2492 |  |  |  |  | Residuals | 24 | 1.0487 | 0.0437 |  |  |  |
| **WEEK4** | log(^α^S_PIE_) | Treatment | 1 | 0.0400 | 0.0400 | 1.197 | 0.2850 |  | log(^α^S_PIE_) | Treatment | 1 | 0.075 | 0.0753 | 0.223 | 0.6410 |  |
|  |  | Treatment:Meta | 4 | 0.0595 | 0.0149 | 0.445 | 0.7750 |  |  | Treatment:Meta | 4 | 0.627 | 0.1567 | 0.464 | 0.7610 |  |
|  |  | Residuals | 24 | 0.8029 | 0.0335 |  |  |  |  | Residuals | 23 | 7.760 | 0.3374 |  |  |  |
| **WEEK 8** | ^α^S_PIE_ | Treatment | 1 | 11.060 | 11.055 | 3.671 | 0.0674 | . | log(^α^S_PIE_) | Treatment | 1 | 0.3868 | 0.3868 | 3.301 | 0.0818 | . |
|  |  | Treatment:Meta | 4 | 2.240 | 0.5600 | 0.186 | 0.9434 |  |  | Treatment:Meta | 4 | 0.2488 | 0.0622 | 0.531 | 0.7143 |  |
|  |  | Residuals | 24 | 72.280 | 3.0120 |  |  |  |  | Residuals | 24 | 2.8122 | 0.1172 |  |  |  |
| **WEEK 12** | ^α^S_PIE_^^0.5^ | Treatment | 1 | 0.020 | 0.0201 | 0.090 | 0.7670 |  | ^α^S_PIE_^^0.5^ | Treatment | 1 | 0.306 | 0.3056 | 0.691 | 0.4140 |  |
|  |  | Treatment:Meta | 4 | 1.780 | 0.445 | 1.990 | 0.1300 |  |  | Treatment:Meta | 4 | 1.012 | 0.2531 | 0.572 | 0.6850 |  |
|  |  | Residuals | 23 | 5.144 | 0.2236 |  |  |  |  | Residuals | 23 | 10.172 | 0.4423 |  |  |  |
| **WEEK 16** | log(^α^S_PIE_) | Treatment | 1 | 0.0084 | 0.0084 | 0.088 | 0.7690 |  | log(^α^S_PIE_) | Treatment | 1 | 2.462 | 2.462 | 7.016 | **0.0141** | ***** |
|  |  | Treatment:Meta | 4 | 0.8831 | 0.2208 | 2.320 | 0.0860 | . |  | Treatment:Meta | 4 | 2.724 | 0.681 | 1.940 | 0.1363 |  |
|  |  | Residuals | 24 | 2.2839 | 0.0952 |  |  |  |  | Residuals | 24 | 8.424 | 0.351 |  |  |  |

Table S13. Summary table of nested ANOVA performed on the number of observed ASVs (^α^S), effective number of ASVs of PIE (^α^S_PIE_) and evenness (J) at α-scale, and Whittaker’s β-diversity (β_S_) in prokaryotes and microeukaryotes. Applied data transformation is presented along with the abbreviation of each variable. Significant P-values are marked in bold. (continued)

|  | **Prokaryotes** | | | | | | | | **Microeukaryotes** | | | | | | | |
| --- | --- | --- | --- | --- | --- | --- | --- | --- | --- | --- | --- | --- | --- | --- | --- | --- |
|  |  |  | Df | Sum Sq | Mean Sq | F-value | P-value |  |  |  | Df | Sum Sq | Mean Sq | F-value | P-value |  |
| **WEEK4** | log(J) | Treatment | 1 | 0.0079 | 0.0079 | 1.640 | 0.2130 |  | J | Treatment | 1 | 0.00175 | 0.0018 | 0.170 | 0.6840 |  |
|  |  | Treatment:Meta | 4 | 0.0086 | 0.0021 | 0.445 | 0.7750 |  |  | Treatment:Meta | 4 | 0.01376 | 0.0034 | 0.334 | 0.8520 |  |
|  |  | Residuals | 24 | 0.1156 | 0.0048 |  |  |  |  | Residuals | 23 | 0.23685 | 0.0103 |  |  |  |
| **WEEK 8** | J^^0.25^ | Treatment | 1 | 0.0007 | 0.0007 | 2.618 | 0.1190 |  | J^^4^ | Treatment | 1 | 0.001596 | 0.0016 | 2.603 | 0.1200 |  |
|  |  | Treatment:Meta | 4 | 0.0006 | 0.0001 | 0.561 | 0.6930 |  |  | Treatment:Meta | 4 | 0.0010 | 0.0003 | 0.423 | 0.7900 |  |
|  |  | Residuals | 24 | 0.0061 | 0.0003 |  |  |  |  | Residuals | 24 | 0.014709 | 0.0006 |  |  |  |
| **WEEK 12** | log(J) | Treatment | 1 | 0.0000 | 0.0000 | 0.005 | 0.9430 |  | J | Treatment | 1 | 0.00448 | 0.0045 | 0.396 | 0.5350 |  |
|  |  | Treatment:Meta | 4 | 0.0335 | 0.0084 | 1.671 | 0.1910 |  |  | Treatment:Meta | 4 | 0.02567 | 0.0064 | 0.568 | 0.6880 |  |
|  |  | Residuals | 23 | 0.1151 | 0.0050 |  |  |  |  | Residuals | 23 | 0.2597 | 0.0113 |  |  |  |
| **WEEK 16** | log(J) | Treatment | 1 | 0.0000 | 0.0000 | 0.000 | 0.9900 |  | log(J) | Treatment | 1 | 0.2751 | 0.2751 | 9.808 | **0.0045** | ** |
|  |  | Treatment:Meta | 4 | 0.0303 | 0.0076 | 1.614 | 0.2030 |  |  | Treatment:Meta | 4 | 0.2009 | 0.0502 | 1.791 | 0.1636 |  |
|  |  | Residuals | 24 | 0.1127 | 0.0047 |  |  |  |  | Residuals | 24 | 0.6731 | 0.0281 |  |  |  |
| **WEEK 4** | β_S_^^0.5^ | Treatment | 1 | 0.0002 | 0.0002 | 0.012 | 0.9130 |  | β_S_ | Treatment | 1 | 0.257 | 0.2569 | 0.188 | 0.6690 |  |
|  |  | Treatment:Meta | 4 | 0.0120 | 0.0030 | 0.173 | 0.9500 |  |  | Treatment:Meta | 4 | 8.992 | 2.2479 | 1.644 | 0.1970 |  |
|  |  | Residuals | 24 | 0.4161 | 0.0173 |  |  |  |  | Residuals | 23 | 31.443 | 1.3671 |  |  |  |
| **WEEK 8** | β_S_ | Treatment | 1 | 0.008 | 0.0076 | 0.019 | 0.8930 |  | log(β_S_) | Treatment | 1 | 0.0010 | 0.0010 | 0.044 | 0.8360 |  |
|  |  | Treatment:Meta | 4 | 0.207 | 0.0517 | 0.127 | 0.9710 |  |  | Treatment:Meta | 4 | 0.0041 | 0.0010 | 0.044 | 0.9960 |  |
|  |  | Residuals | 24 | 9.756 | 0.4065 |  |  |  |  | Residuals | 24 | 0.5607 | 0.0234 |  |  |  |
| **WEEK 12** | β_S_ | Treatment | 1 | 1.052 | 1.0517 | 4.284 | **0.0499** | ***** | log(β_S_) | Treatment | 1 | 0.0276 | 0.0276 | 1.053 | 0.3160 |  |
|  |  | Treatment:Meta | 4 | 0.430 | 0.1075 | 0.438 | 0.7798 |  |  | Treatment:Meta | 4 | 0.1125 | 0.0281 | 1.072 | 0.3930 |  |
|  |  | Residuals | 23 | 5.647 | 0.2455 |  |  |  |  | Residuals | 23 | 0.6037 | 0.0263 |  |  |  |
| **WEEK 16** | β_S_ | Treatment | 1 | 0.081 | 0.0806 | 0.2290 | 0.6370 |  | β_S_ | Treatment | 1 | 0.2700 | 0.2695 | 0.270 | 0.6080 |  |
|  |  | Treatment:Meta | 4 | 0.226 | 0.0565 | 0.1600 | 0.9560 |  |  | Treatment:Meta | 4 | 0.1140 | 0.0284 | 0.028 | 0.9980 |  |
|  |  | Residuals | 24 | 8.462 | 0.3526 |  |  |  |  | Residuals | 24 | 23.971 | 0.9988 |  |  |  |

Table S14. Summary table of generalized additive models (GAMs) performed on the number of observed ASVs (^α^S) in prokaryotes and microeukaryotes over the entire experimental duration. The models included treatment as the main linear predictor and time (i.e., the week of sampling) with varying shapes of smooth according to individual metacommunities (k=3 and k=4 for prokaryotes and microeukaryotes, respectively). Applied data transformation is presented along with the abbreviation of each variable. Significant P-values are marked in bold.

|  | **Prokaryotes** | | | | | |
| --- | --- | --- | --- | --- | --- | --- |
|  |  | Estimate | Std. Error | t-value | P-value |  |
| ^α^S | Intercept | 246.577 | 5.475 | 45.036 | <2e-16 | *** |
|  | Treatment[fragm.] | -11.831 | 7.710 | -1.534 | 0.1280 |  |
|  |  | edf | Ref. df | F-value | P-value |  |
|  | s(week):M1 | 1.709 | 1.916 | 1.143 | 0.3109 |  |
|  | s(week):M2 | 1.510 | 1.760 | 0.522 | 0.6145 |  |
|  | s(week):M3 | 1.000 | 1.000 | 1.779 | 0.1851 |  |
|  | s(week):M4 | 1.000 | 1.001 | 1.784 | 0.1844 |  |
|  | s(week):M5 | 1.765 | 1.945 | 5.185 | **0.0176** | ***** |
|  | s(week):M6 | 1.758 | 1.941 | 3.246 | 0.0747 | . |
|  | R^2^_adj_ | 0.125 |  |  |  |  |
|  | AIC | 1241.431 |  |  |  |  |
|  | **Microeukaryotes** | | | | | |
|  |  | Estimate | Std. Error | t-value | P-value |  |
| log(^α^S) | Intercept | 4.952 | 0.028 | 175.377 | <2e-16 | *** |
|  | Treatment[fragm.] | -0.103 | 0.040 | -2.601 | **0.0107** | ***** |
|  |  | edf | Ref. df | F-value | P-value |  |
|  | s(week):M1 | 2.396 | 2.744 | 4.618 | **0.0040** | ****** |
|  | s(week):M2 | 2.507 | 2.822 | 4.662 | **0.0032** | ****** |
|  | s(week):M3 | 2.783 | 2.963 | 13.447 | **2.41e-07** | ******* |
|  | s(week):M4 | 1.857 | 2.235 | 0.747 | 0.4103 |  |
|  | s(week):M5 | 2.686 | 2.925 | 3.638 | **0.0093** | ****** |
|  | s(week):M6 | 2.021 | 2.404 | 0.935 | 0.3021 |  |
|  | R^2^_adj_ | 0.441 |  |  |  |  |
|  | AIC | -7.675 |  |  |  |  |

Table S15. Summary table of nested ANOVA performed on chlorophyll-a fluorescence data (Chl-a, a.u.). Applied data transformation is presented.

|  |  |  | Df | Sum Sq | Mean Sq | F-value | P-value |
| --- | --- | --- | --- | --- | --- | --- | --- |
| **WEEK 4** | Chl-a^^0.333^ | Treatment | 1 | 0.1160 | 0.1156 | 0.124 | 0.7280 |
|  |  | Treatment:Meta | 4 | 5.3090 | 1.3273 | 1.423 | 0.2570 |
|  |  | Residuals | 21 | 22.3790 | 0.9324 |  |  |
| **WEEK 8** | log(Chl-a) | Treatment | 1 | 0.4620 | 0.4621 | 0.572 | 0.4570 |
|  |  | Treatment:Meta | 4 | 2.1810 | 0.5452 | 0.674 | 0.6160 |
|  |  | Residuals | 21 | 19.4030 | 0.8085 |  |  |
| **WEEK 12** | log(Chl-a) | Treatment | 1 | 0.1980 | 1.0587 | 0.283 | 0.5990 |
|  |  | Treatment:Meta | 4 | 4.2350 | 0.0667 | 1.516 | 0.2290 |
|  |  | Residuals | 24 | 16.7620 | 0.6984 |  |  |
| **WEEK 16** | log(Chl-a) | Treatment | 1 | 0.4800 | 0.4803 | 1.055 | 0.3150 |
|  |  | Treatment:Meta | 4 | 1.5410 | 0.3853 | 0.847 | 0.5100 |
|  |  | Residuals | 24 | 10.9210 | 0.4551 |  |  |

Table S16. Summary table of nested ANOVA performed on the number of observed species (^α^S), the effective number of species of PIE (^α^S_PIE_) and evenness (J) at α-scale in phytoplankton in week 16. Applied data transformation is presented along with the abbreviation of each variable. Significant P-values are marked in bold.

|  |  | Df | Sum Sq | Mean Sq | F-value | P-value |  |
| --- | --- | --- | --- | --- | --- | --- | --- |
| log(^α^S) | Treatment | 1 | 0.0277 | 0.0277 | 0.328 | 0.5720 |  |
|  | Treatment:Meta | 4 | 0.7079 | 0.1770 | 2.097 | 0.1130 |  |
|  | Residuals | 24 | 2.0256 | 0.0844 |  |  |  |
| ^α^S_PIE_^^0.25^ | Treatment | 1 | 0.2756 | 0.2756 | 6.226 | **0.0199** | ***** |
|  | Treatment:Meta | 4 | 0.1159 | 0.0290 | 0.654 | 0.6294 |  |
|  | Residuals | 24 | 1.0624 | 0.0443 |  |  |  |
| asin(J^^0.5^) | Treatment | 1 | 0.2589 | 0.2589 | 3.958 | 0.0582 | . |
|  | Treatment:Meta | 4 | 0.1314 | 0.0328 | 0.502 | 0.7345 |  |
|  | Residuals | 24 | 1.5698 | 0.0654 |  |  |  |

Table S17. ANOVA table for best fit linear mixed-effects models (LMMs) demonstrating the effect of the treatment and zooplankton biomass (ZB) on the number of observed ASVs (^α^S), effective number of ASVs of PIE (^α^S_PIE_) and evenness (J) in prokaryotes and microeukaryotes on week 16. In the LMMs, treatment and ZB were included as fixed, and metacommunity ID as a nested random effect factor. Applied data transformation is presented along with the abbreviation of each variable. Significant P-values are marked in bold.

|  | **Prokaryotes** | | | | | |
| --- | --- | --- | --- | --- | --- | --- |
|  |  | F | Df | Df. res | P-value |  |
| ^α^S | Treatment | 0.9368 | 1 | 4.7903 | 0.3794 |  |
|  | log(ZB) | 0.0018 | 1 | 23.1829 | 0.9662 |  |
| log(^α^S_PIE_) | Treatment | 0.1694 | 1 | 4.5008 | 0.6995 |  |
|  | log(ZB) | 0.5549 | 1 | 26.8518 | 0.4628 |  |
| log(J) | Treatment | 0.1105 | 1 | 4.6274 | 0.7541 |  |
|  | log(ZB) | 0.8001 | 1 | 26.9056 | 0.3790 |  |
|  | **Microeukaryotes** | | | | | |
|  |  | F | Df | Df. res | P-value |  |
| log(^α^S) | Treatment | 11.4515 | 1 | 4.5444 | **0.02272** | ***** |
|  | log(ZB) | 4.1106 | 1 | 24.1278 | 0.05380 | . |
|  | Treatment*log(ZB) | 7.1437 | 1 | 25.4269 | **0.01295** | * |
| log(^α^S_PIE_) | Treatment | 1.0506 | 1 | 4.7903 | 0.354323 |  |
|  | log(ZB) | 10.7387 | 1 | 23.1829 | **0.003285** | ** |
| log(J) | Treatment | 2.4915 | 1 | 4.7903 | 0.177827 |  |
|  | log(ZB) | 9.0677 | 1 | 23.1829 | **0.006189** | ** |

Table S18. Summary table of the structural equation model (SEM) representing the relationships among treatment, the number of observed microeukaryote ASVs (^α^S), zooplankton biomass (ZB) and evenness (J) in microeukaryotes in week 16. Applied data transformation is presented along with the abbreviation of each variable. In the models, microeukaryote log(^α^S), log(ZB) and treatment were included as fixed, and metacommunity ID as a nested random effect factor. Effects for which the confidence intervals do not contain zero are highlighted with a star (i.e., significant at the CI level).

|  |  |  | Std. Effect | Bias | Std. Error | CI (95%) |  |
| --- | --- | --- | --- | --- | --- | --- | --- |
| log(^α^S) | Direct | Treatment[fragm.] | -0.612 | 0.018 | 0.119 | -0.859 – -0.404 | ***** |
|  | Total | Treatment[fragm.] | -0.612 | 0.018 | 0.119 | -0.859 – -0.404 | ***** |
| log(J) | Direct | Treatment[fragm.] | -0.228 | -0.018 | 0.086 | -0.415 – -0.078 | ***** |
|  |  | log(ZB) | 0.476 | -0.063 | 0.149 | 0.201 – 0.674 | ***** |
|  | Indirect | Treatment[fragm.] | -0.167 | 0.030 | 0.060 | -0.273 – -0.055 | ***** |
|  |  | log(^α^S) | 0.173 | -0.064 | 0.101 | 0.036 – 0.464 | ***** |
|  | Total | Treatment[fragm.] | -0.394 | 0.012 | 0.084 | -0.538 – -0.229 | ***** |
|  |  | log(^α^S) | 0.173 | -0.064 | 0.101 | 0.036 – 0.464 | ***** |
|  |  | log(ZB) | 0.476 | -0.063 | 0.149 | 0.201 – 0.674 | ***** |
|  | Mediators | log(^α^S) | -0.106 | 0.044 | 0.055 | -0.259 – -0.028 | ***** |
|  |  | log(ZB) | 0.006 | -0.036 | 0.066 | -0.063 – 0.219 |  |
| log(ZB) | Direct | Treatment[fragm.] | -0.128 | -0.083 | 0.120 | -0.279 – 0.045 |  |
|  |  | log(^α^S) | 0.363 | -0.156 | 0.202 | 0.151 – 0.661 | ***** |
|  | Indirect | Treatment[fragm.] | -0.222 | 0.100 | 0.109 | -0.376 – -0.122 | ***** |
|  | Total | Treatment[fragm.] | 0.350 | 0.016 | 0.112 | -0.548 – -0.093 | ***** |
|  |  | log(^α^S) | 0.363 | -0.156 | 0.202 | 0.151 – 0.661 | ***** |
|  | Mediators | log(^α^S) | -0.222 | 0.100 | 0.109 | -0.376 – -0.122 | ***** |

Table S19. Fixed effects table of the generalized linear mixed-effects models (GLMMs) performed on the presence/absence of the ASVs in week 16 in prokaryotes and microeukaryotes. In the models, regional abundance in week 4 (RA), zooplankton biomass in week 16 (ZB), and treatment were included as fixed, and metacommunity ID as a nested random effect factor.

|  | **Prokaryotes** | | | | | |
| --- | --- | --- | --- | --- | --- | --- |
|  |  | Odds ratio | CI (95%) | z-value | P-value |  |
| Presence in week 16 | Intercept | 0.09 | 0.08 – 0.09 | -69.444 | <2e-16 | ******* |
|  | RA | 1.24 | 1.14 – 1.36 | 4.961 | **7.01e-07** | ******* |
|  | ZB | 1.05 | 1.00 – 1.11 | 1.888 | 0.0591 | **.** |
|  | Treatment[fragm.] | 1.44 | 0.95 – 2.18 | 1.723 | 0.0849 | **.** |
|  | RA*Treatment[fragm.] | 1.64 | 1.30 – 2.06 | 4.230 | **2.33e-05** | ******* |
|  | ZB*Treatment[fragm.] | 1.87 | 0.89 – 3.95 | 1.650 | 0.0990 |  |
|  | **Microeukaryotes** | | | | | |
|  |  | Odds ratio | CI (95%) | z-value | P-value |  |
| Presence in week 16 | Intercept | 0.09 | 0.08 – 0.09 | -108.026 | <2e-16 | *** |
|  | RA | 1.02 | 1.00 – 1.05 | 2.155 | **0.0312** | ***** |
|  | ZB | 1.21 | 1.18 – 1.25 | 12.502 | **<2e-16** | ******* |
|  | Treatment[fragm.] | 0.59 | 0.46 – 0.75 | -4.289 | **7.67e-06** | ******* |
|  | RA*Treatment[fragm.] | 1.02 | 0.99 – 1.06 | 1.282 | 0.1999 |  |
|  | ZB*Treatment [fragm.] | 0.55 | 0.36 – 0.84 | -2.756 | **0.0059** | ****** |

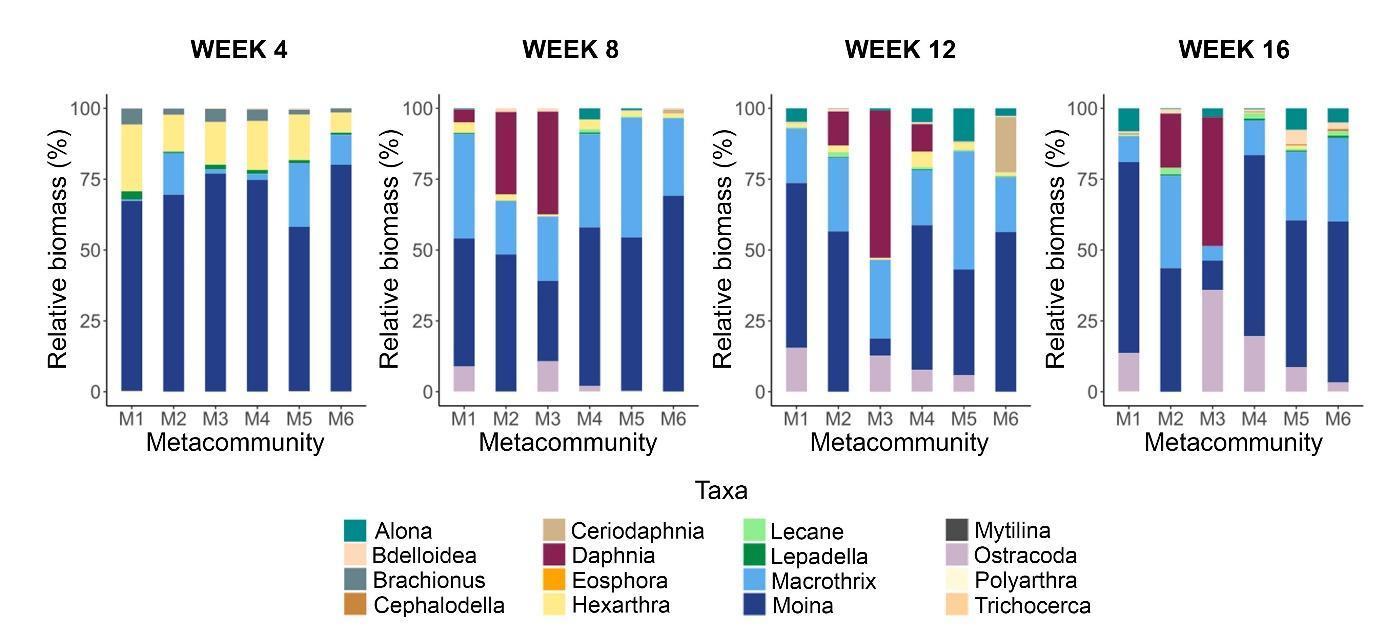
Figure S1. Relative biomasses (%) of zooplankton taxa over the experimental period separately shown for each metacommunity (i.e, mean values of 5 mesocosms). Treatment assignment: metacommunity M1-M3: connectivity, metacommunity M4-M6: fragmentation.

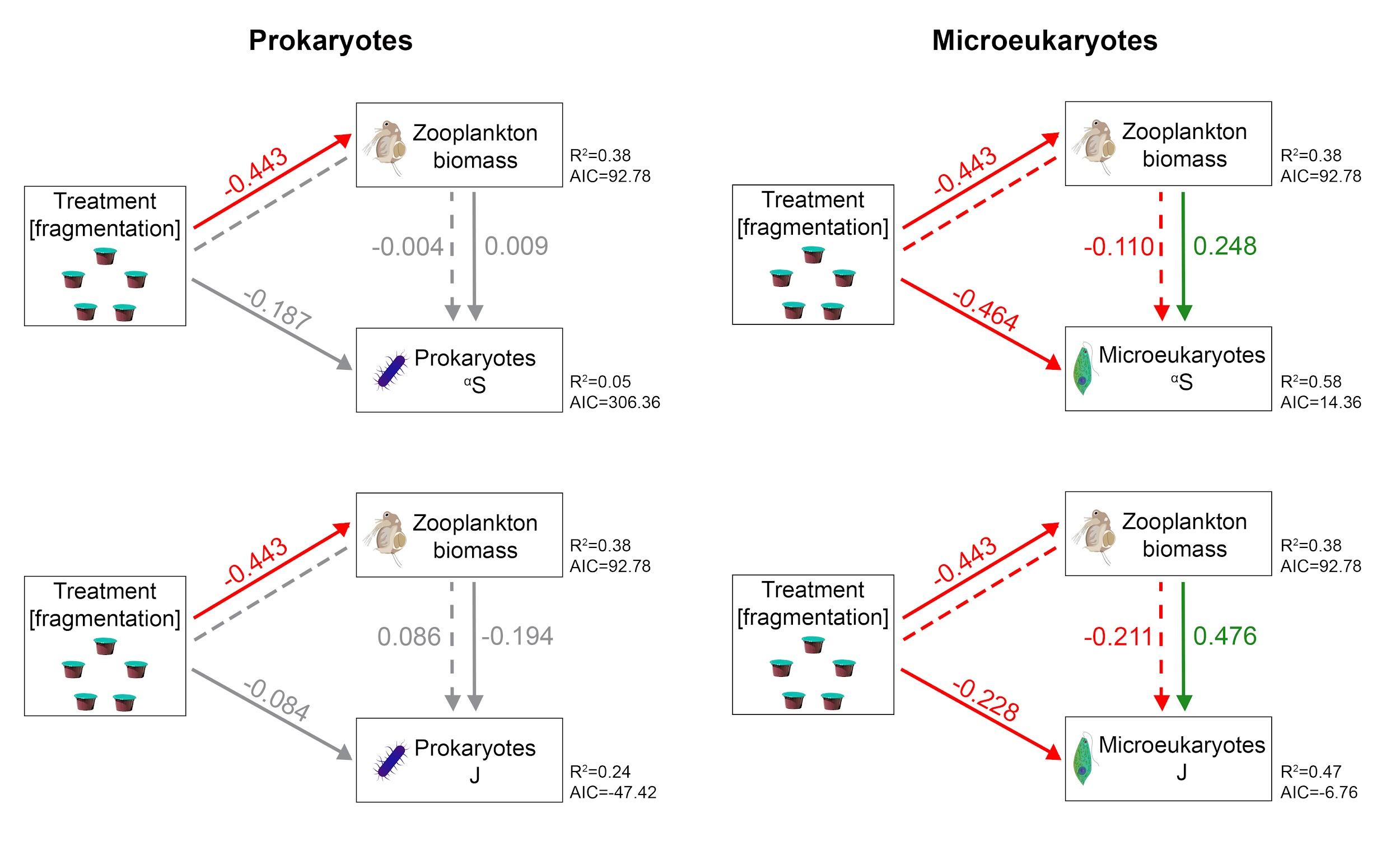
Figure S2. Structural equation models (SEMs) representing the relationships among treatment, zooplankton biomass, the number of observed ASVs (^α^S) and evenness (J) in prokaryotes and microeukaryotes in week 16. Variables are represented by boxes and the directional relationships by single-head arrows (solid line: direct effect; dashed line: indirect effect; green: significant positive effect; grey: non-significant effect). Standardized coefficients are shown on the side of the arrows. R^2^ and AIC values for the component models are indicated next to the boxes of endogenous variables. Zooplankton biomass and J were log-transformed in each case, and ^α^S in the case of microeukaryotes. In the models, treatment and zooplankton biomass were included as fixed, and metacommunity ID as a nested random effect factor.

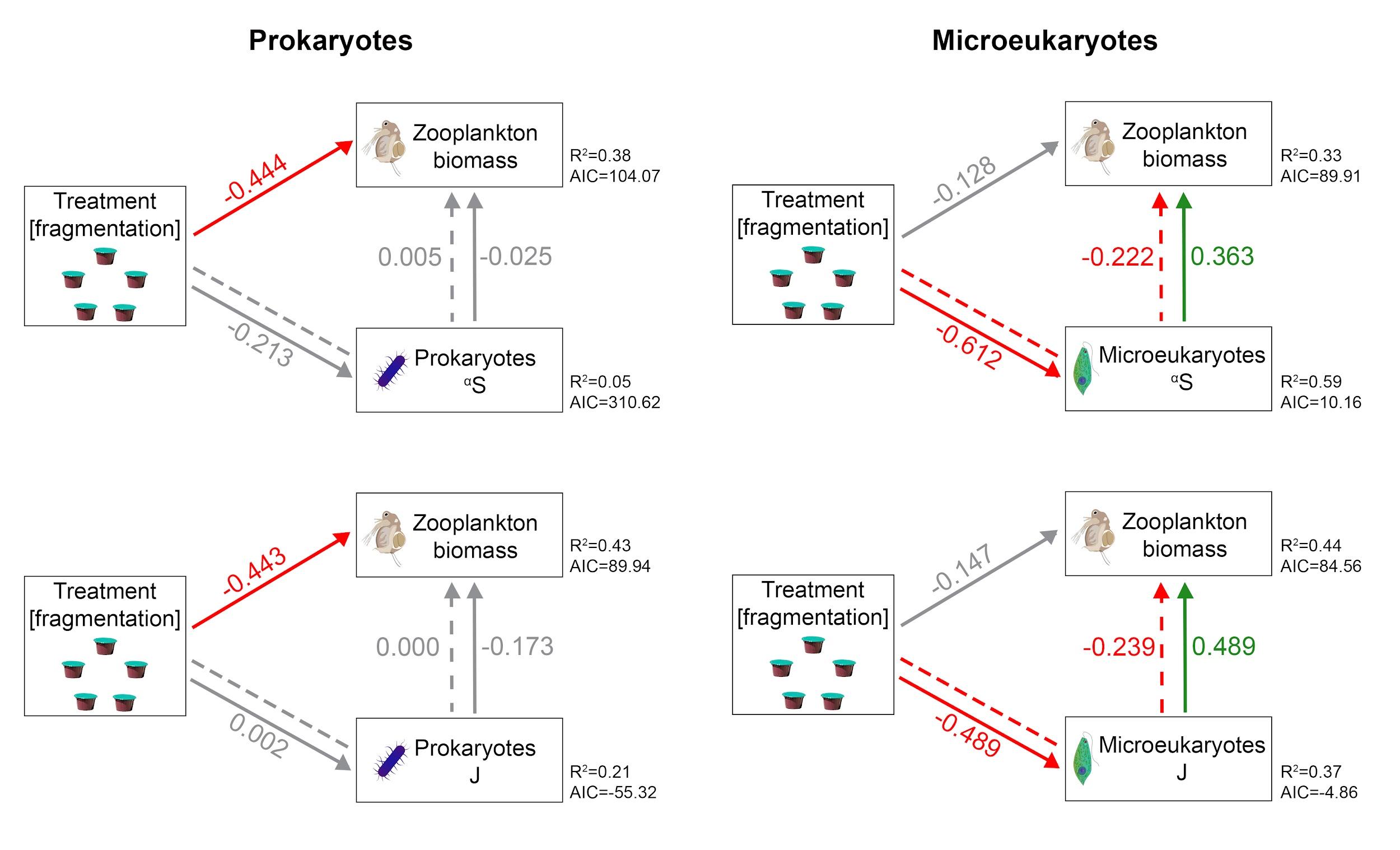
Figure S3. Structural equation models (SEMs) representing the relationships among treatment, zooplankton biomass (ZB), the number of observed ASVs (^α^S) and evenness (J) in prokaryotes and microeukaryotes in week 16. Variables are represented by boxes and the directional relationships by single-head arrows (solid line: direct effect; dashed line: indirect effect; green: significant positive effect; grey: non-significant effect). Standardized coefficients are shown on the side of the arrows. R^2^ and AIC values for the component models are indicated next to the boxes of endogenous variables. Zooplankton biomass and J were log-transformed in each case, and ^α^S in the case of microeukaryotes. In the models, treatment and zooplankton biomass were included as fixed, and metacommunity ID as a nested random effect factor.

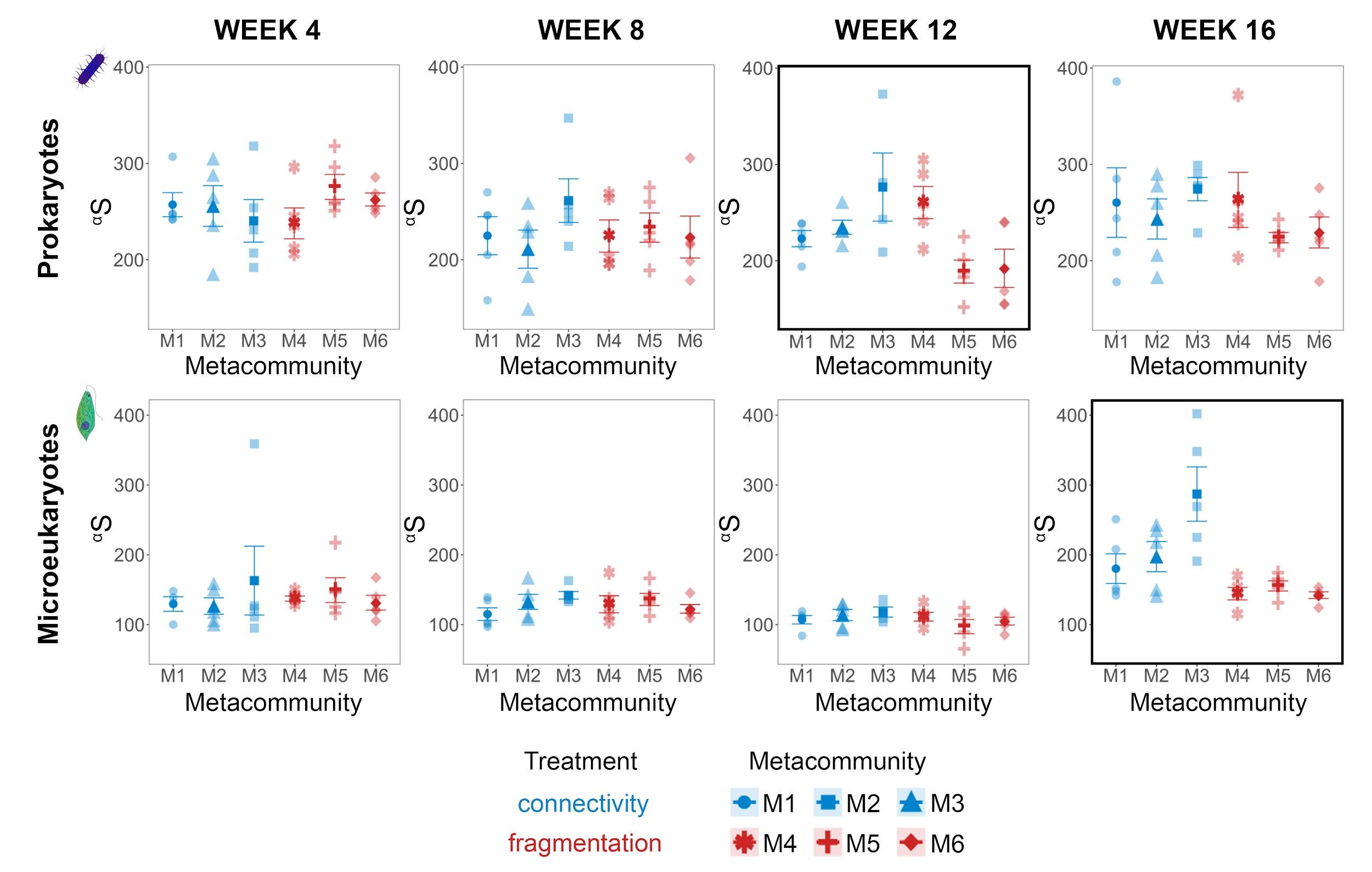
Figure S4. Number of observed ASVs at α-scale (^α^S) of prokaryotes and microeukaryotes in each metacommunity shown separately for each sampling date. The boxes marked with a bold black frame indicate a significant treatment effect (P<0.05) resulting from the nested ANOVA.

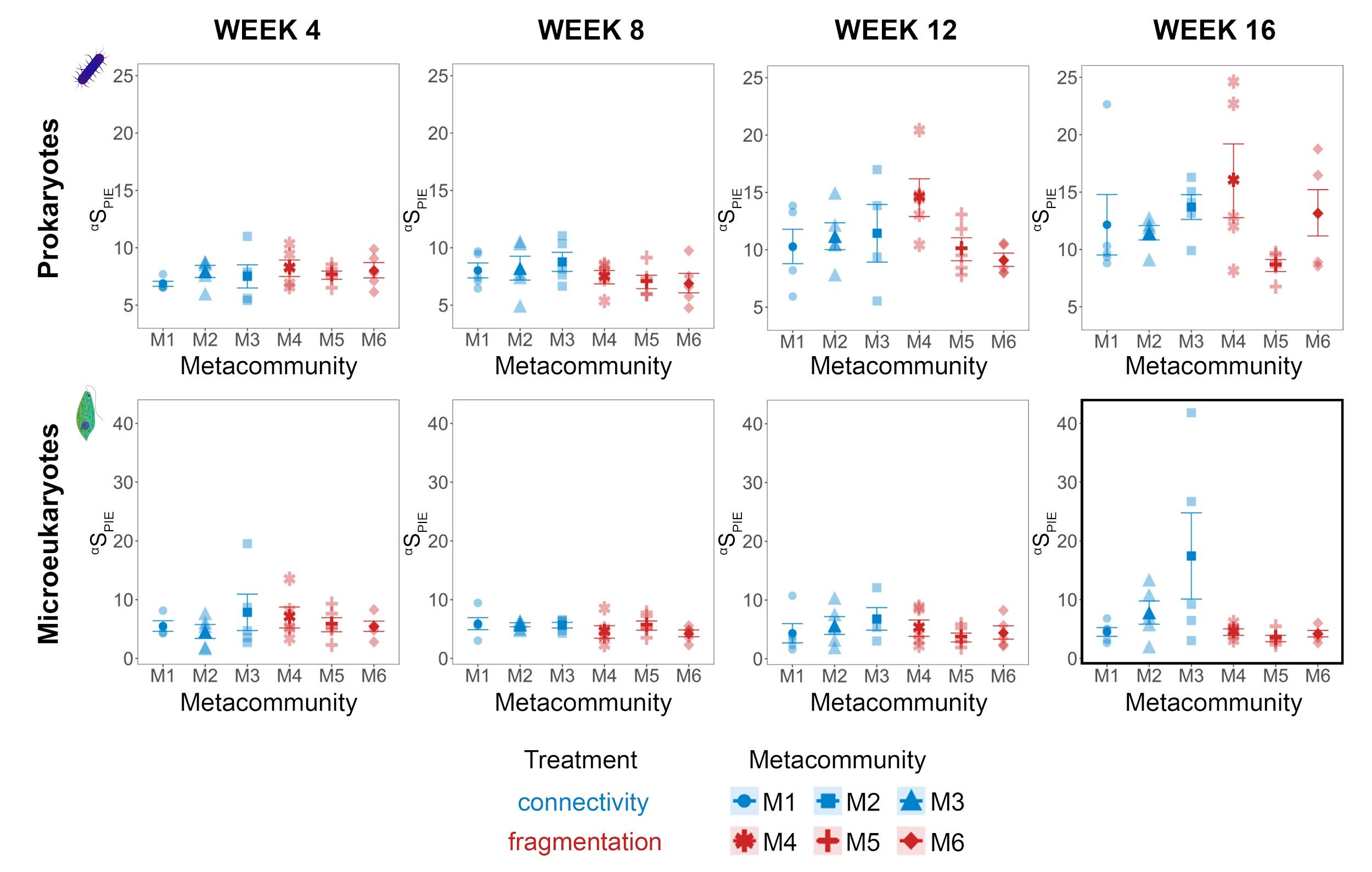
Figure S5. Effective number of ASVs of PIE at α-scale (^α^S_PIE_) of prokaryotes and microeukaryotes in each metacommunity shown separately for each sampling date. The box marked with a bold black frame indicates a significant treatment effect (P<0.05) resulting from the nested ANOVA.

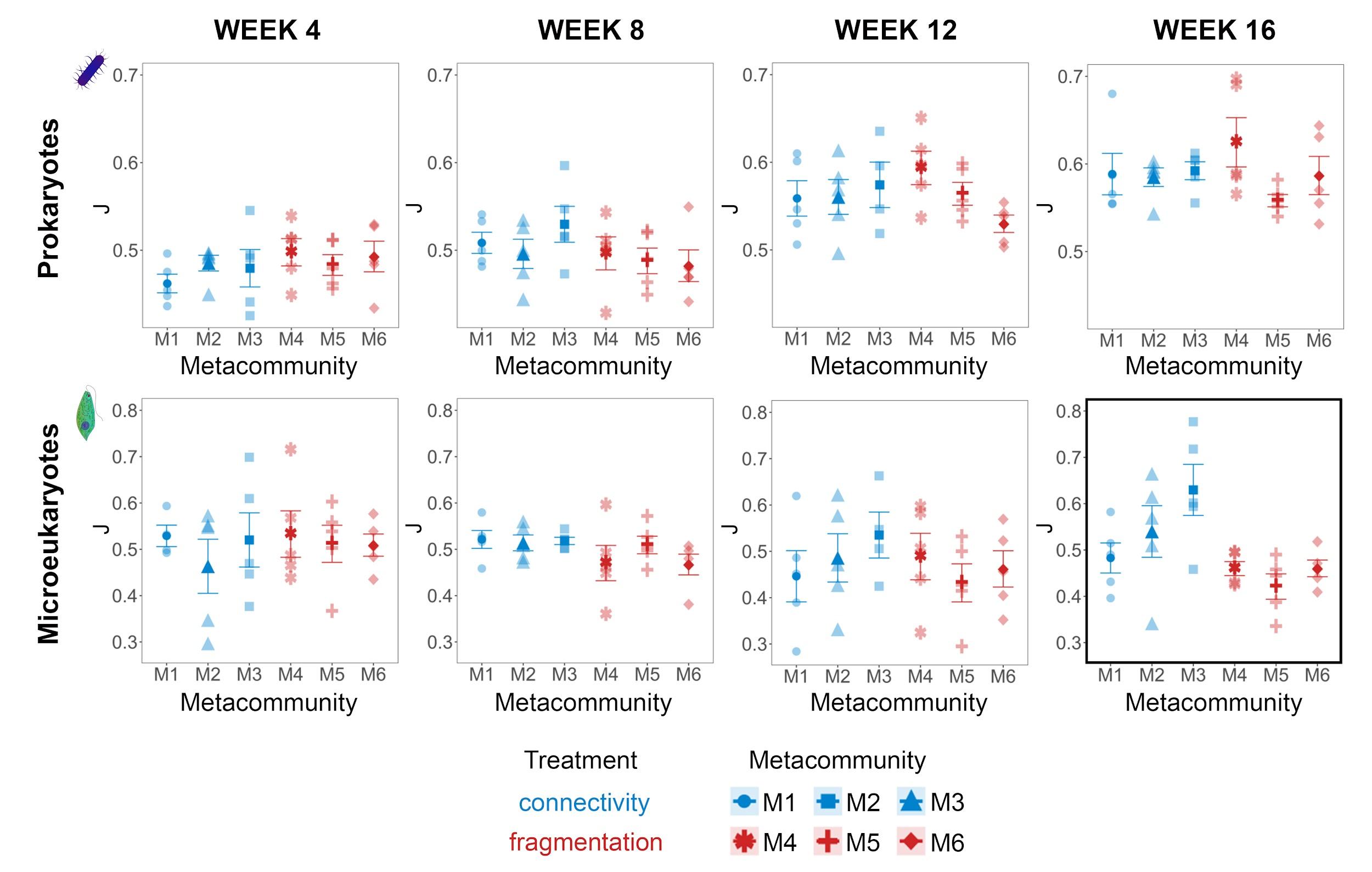
Figure S6. Evenness (J) of prokaryotes and microeukaryotes in each metacommunity shown separately for each sampling date. The box marked with a bold black frame indicates a significant treatment effect (P<0.05) resulting from the nested ANOVA.

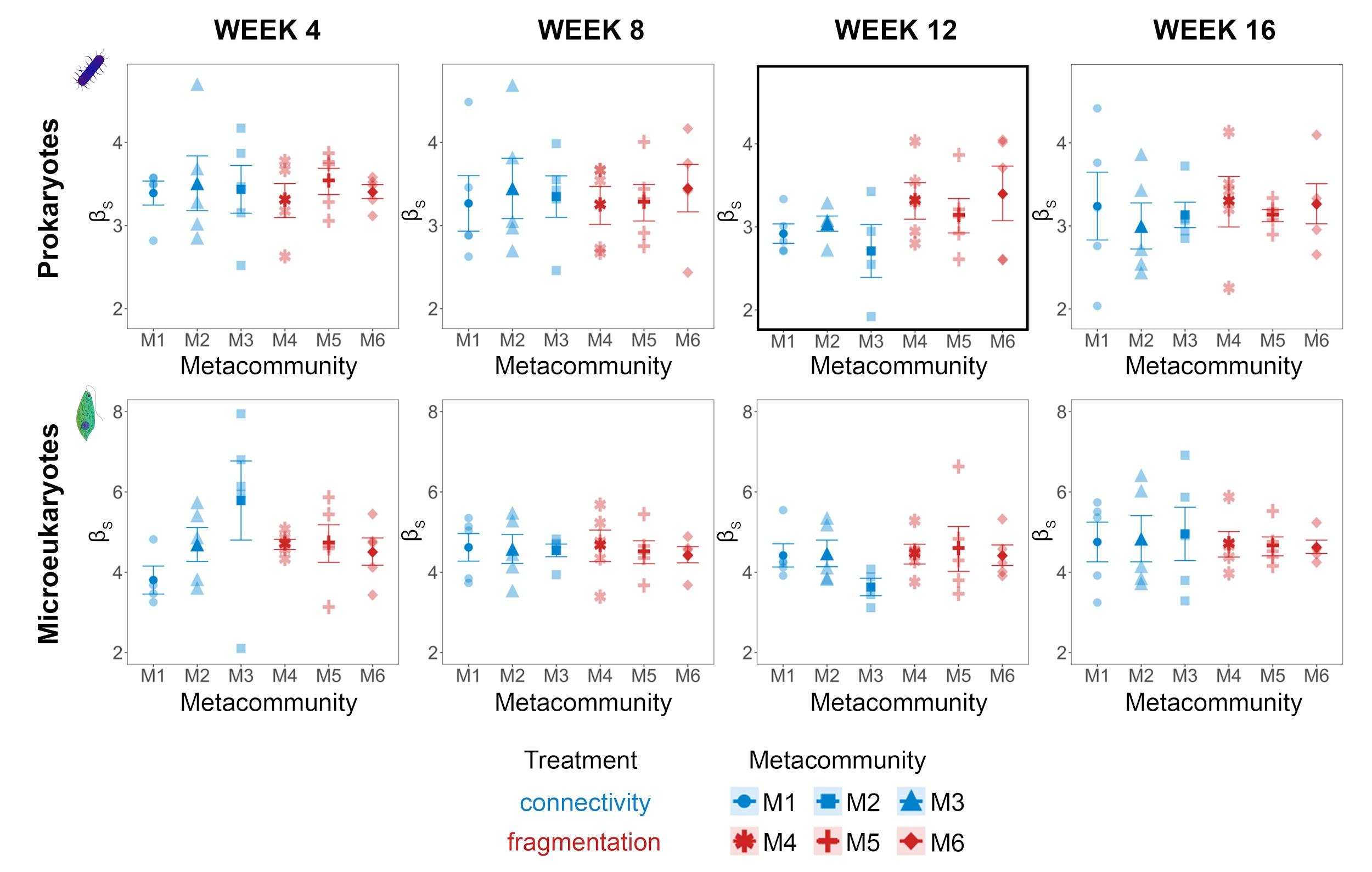
Figure S7. Whittaker’s β-diversity (β_S_) of prokaryotes and microeukaryotes in each metacommunity shown separately for each sampling date. The box marked with a bold black frame indicates a significant treatment effect (P<0.05) resulting from the nested ANOVA.

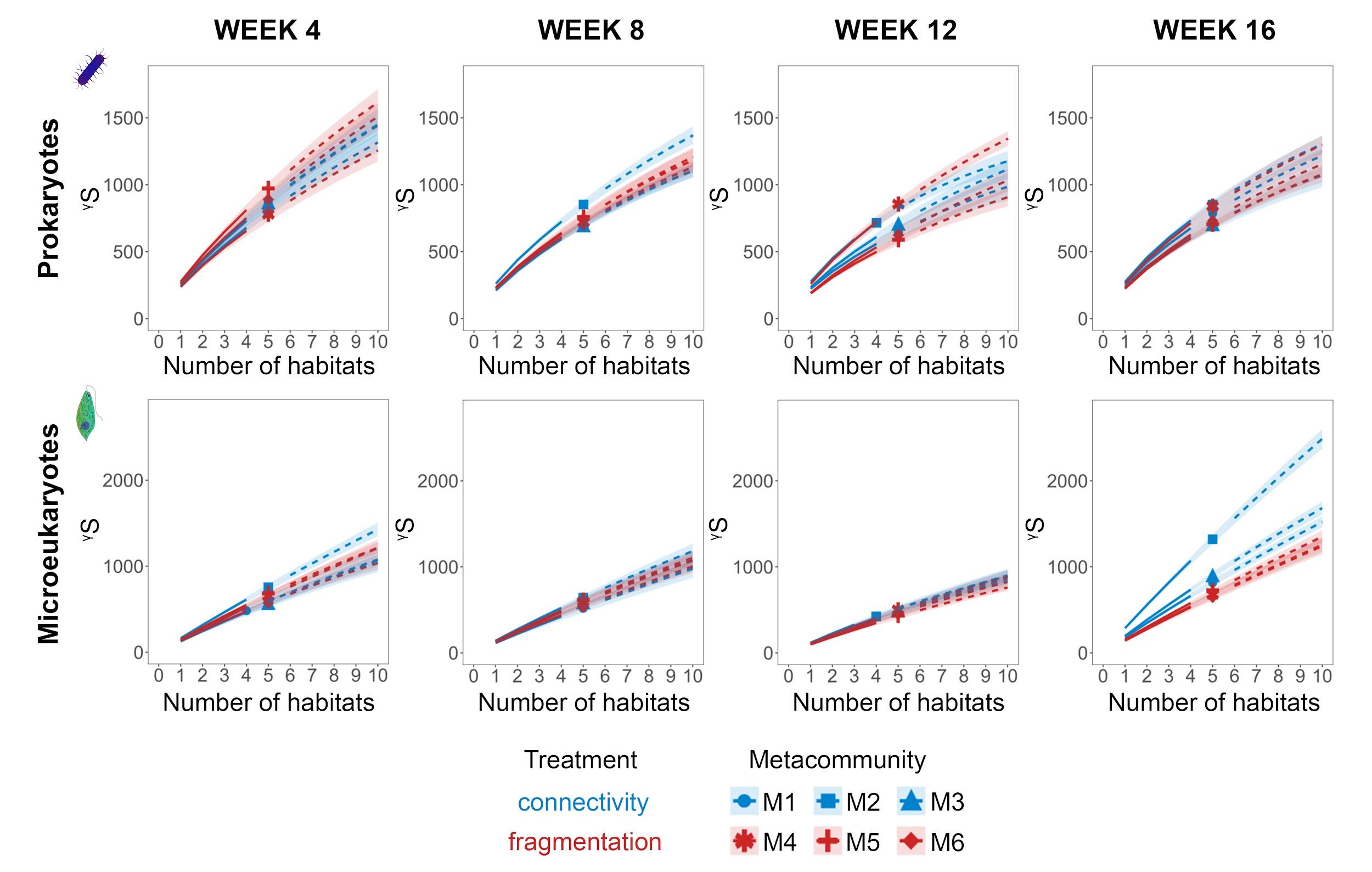
Figure S8. Accumulation curves with the number of observed ASVs at γ-scale (^γ^S), extrapolated estimates (dashed lines), and 95% confidence intervals (error bands) for prokaryotes and microeukaryotes in each metacommunity shown separately for each sampling date.

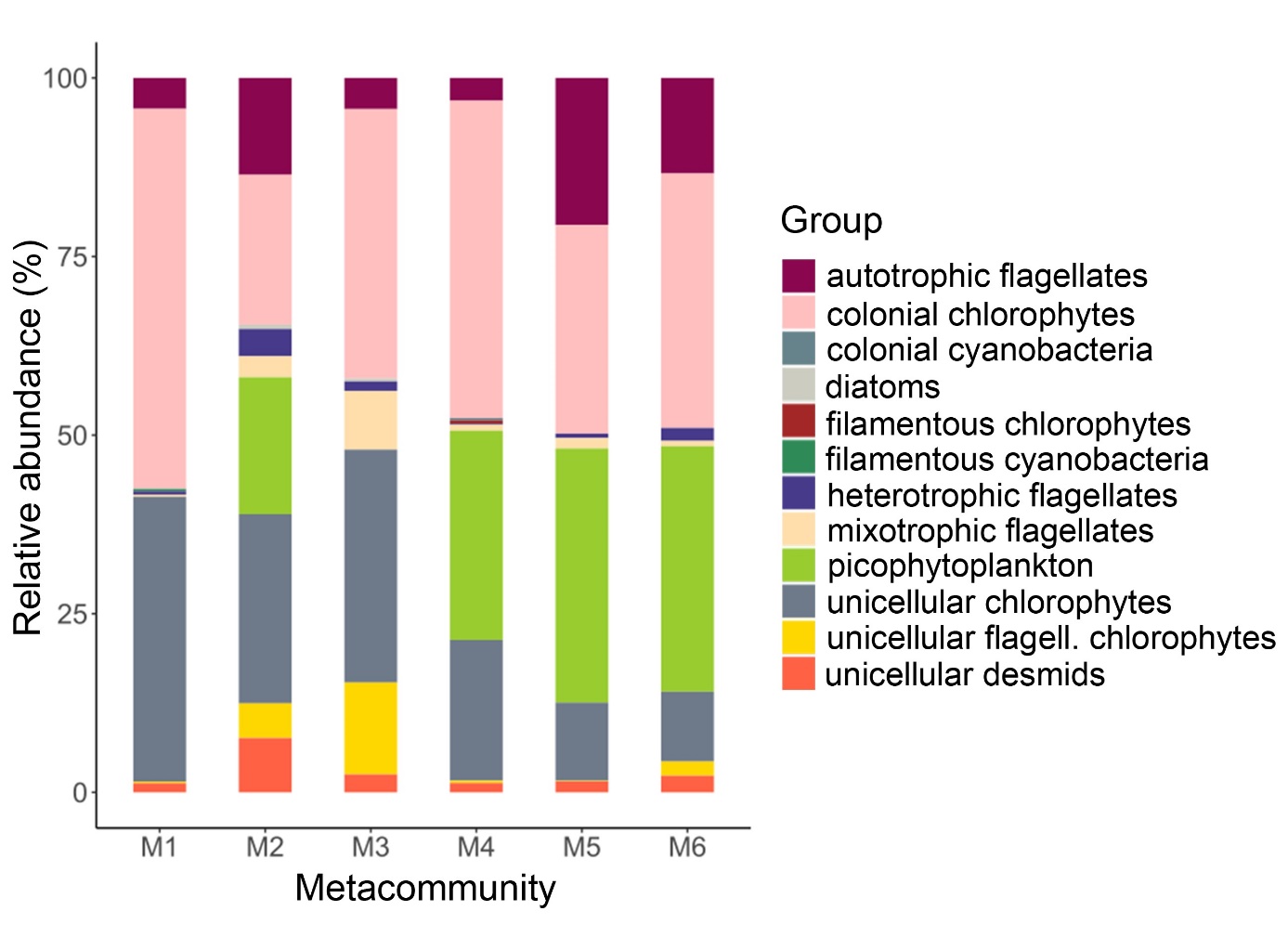

Figure S9. Relative abundances (%) of groups of phytoplankton taxa in week 16 separately shown for each metacommunity (i.e, mean values of 5 mesocosms). Treatment assignment: metacommunity M1-M3: connectivity, metacommunity M4-M6: fragmentation. Cell count data (cells mL^-1^) of phytoplankton taxa are presented in Table S20.
